# Silencing *SmD1* in Solanaceae alters susceptibility to root-knot nematodes

**DOI:** 10.1101/2020.11.25.398149

**Authors:** Joffrey Mejias, Yongpan Chen, Nhat-My Truong, Karine Mulet, Stéphanie Jaubert-Possamai, Pierre Abad, Bruno Favery, Michaël Quentin

## Abstract

Root-knot nematodes (RKNs) are among the most damaging pests of agricultural crops. Indeed, *Meloidogyne* is an extremely polyphagous genus of nematodes that can infect thousands of plant species. A few genes for resistance (R-genes) to RKNs suitable for use in crop breeding have been identified, and new virulent strains and species of nematode emerge rendering these R-genes ineffective. Effective parasitism is dependent on the secretion, by the RKN, of effectors targeting plant functions, which mediate the reprogramming of root cells into specialised feeding cells. These cells, the giant cells, are essential for RKN development and reproduction. The EFFECTOR 18 protein (EFF18) from *M. incognita* interacts with the spliceosomal protein SmD1 in Arabidopsis, disrupting its function in alternative splicing regulation and modulating the giant cell transcriptome. We show here that EFF18 is a conserved RKN-specific effector. We also show here that EFF18 effectors also target SmD1 in *Nicotiana benthamiana* and *Solanum lycopersicum*. The alteration of *SmD1* expression by virus-induced gene silencing (VIGS) in Solanaceae affects giant cell formation and nematode development. Thus, *SmD1* is a susceptibility gene and a promising target for the development of broad resistance, especially in Solanaceae, for the control of *Meloidogyne* spp.

## Introduction

Plant parasitic nematodes are major crop pests causing crop losses of several million dollars annually, through damage to almost all cultivated plants, and the transmission of plant viruses (Singh et al., 2013). Root-knot nematodes (RKNs) of the genus *Meloidogyne* are considered to be the most detrimental of these plant parasites, due to the magnitude of the economic losses they cause (Jones *et al.*, 2013). RKNs are widespread worldwide and can infect more than 5,500 different plant species, including many species of major agricultural interest. About 100 RKN species have been described, and those reproducing asexually by mitotic parthenogenesis (*M. incognita*, *M. javanica*, *M. arenaria* and *M. enterolobii*) are the most polyphagous and damaging pests. By contrast, those reproducing sexually or by meiotic parthenogenesis (*M. hapla*) have a smaller host range (Blok et al., 2008; Castagnone-Sereno, 2006).

All RKNs are sedentary endoparasites that induce the formation of specialised feeding structures and typical root deformations, known as galls or root knots, that deprive the plant of nutrients (Escobar *et al.*, 2015; Favery *et al.*, 2016). After hatching from eggs, the stage 2 juveniles (J2) of *M. incognita* penetrate the root apex and migrate between plant cells to reach the plant vascular system (Holbein *et al.*, 2019). Once there, the filiform J2 switch to a sedentary lifestyle, by selecting five to seven cells of the vascular parenchyma and inducing their reprogramming into specialised feeding cells, known as giant cells (Escobar *et al.*, 2015; Favery *et al.*, 2016; Olmo *et al.*, 2020). These hypertrophied and multinucleate cells act as metabolic sinks close to the xylem and phloem vessels that withdraw water and nutrients from the sap (Rodiuc *et al.*, 2014). The nematode uses these specific giant cells for feeding for the rest of its life. After successive moults, the sedentary swollen juveniles develop into an adult female that lays her egg masses on the root surface, thus completing the cycle. The giant cells are hypertrophied and multinucleate, harbouring hundreds of nuclei. They are produced by successive nuclear divisions uncoupled from cytokinesis, followed by nuclear endoreduplication (de Almeida Engler and Gheysen, 2013). RKN induce giant cells and gall formation by recruiting the developmental pathways of post-embryonic organogenesis and regeneration to promote transient pluripotency (Olmo *et al.*, 2020).

RKNs parasitise plants and induce the redifferentiation of vascular cells into giant cells by secreting effectors, molecules that recruit/hijack plant functions (Mejias *et al.*, 2019; Toruño *et al.*, 2016). RKN effectors, particularly those produced by the three oesophageal gland cells and secreted into the host through a stylet, are involved in the four main functions underlying parasitism: (i) the degradation and modification of plant cell walls during J2 migration within the root; (ii) the suppression of host defences; (iii) the reprogramming of plant vascular cells as giant cells and (vi) the maintenance of these feeding sites (Mitchum et al., 2013; Truong et al., 2015). The profound morphological and metabolic changes associated with giant cell induction by RKNs and the transcriptional reprogramming occurring during the formation of these cells require the secretion of effectors targeting key nuclear functions (Hewezi and Baum, 2013; Quentin *et al.*, 2013). With the exception of plant cell wall-degrading enzymes (Danchin *et al.*, 2010), very few effectors have been shown to be conserved and functional in multiple RKN species. For example, 16D10 encodes a conserved secretory peptide conserved in five RKN species (*M. incognita*, *M. arenaria*, *M. hapla*, *M. javanica*, *M. chitwoodi*) that stimulates root growth and functions as a ligand for a putative plant transcription factor (Huang et al., 2006; Dinh, 2015). The silencing of *16D10* by RNA interference methods confers broad resistance to RKNs (Huang et al., 2006; Dinh, 2015). The chorismate mutates, MiCM3 (Wang *et al.*, 2018) and MjCM1 (Doyle and Lambert, 2003), and the transthyretin-like proteins, MjTTL5 (Lin *et al.*, 2016) and MhTTL2 (Gleason *et al.*, 2017), also appear to be effectors conserved among RKNs. Interestingly, MhTTL2 is expressed in the amphids (Gleason *et al.*, 2017), whereas MjTTL5 is expressed specifically in the subventral glands, suggesting different roles for these two molecules in parasitism, encoded by the same gene family (Lin *et al.*, 2016).

We recently showed that MiEFF18, a nuclear effector from *M. incognita,* is secreted *in planta*, targets the giant cell nuclei and interacts with the SmD1 protein, a core component of the spliceosome (Mejias *et al.*, 2020). We show here that MiEFF18 is a specific and conserved RKN effector and that orthologous genes are specifically expressed in the salivary glands of RKNs. We also show that MiEFF18 and its orthologue in *M. enterolobii,* MeEFF18, interact with SmD1 proteins from different plant species. Moreover, virus-induced gene silencing (VIGS) approaches silencing the SmD1 genes of *N. benthamiana* and *S. lycopersicum* greatly impair RKN infection. These results are consistent with the targeting, by RKNs, of conserved spliceosomal functions, to drive the development of giant cells, facilitating parasitism on a large spectrum of host plants.

## Results

### EFF18 is a conserved RKN-specific effector targeting plant nucleus

MiEFF18 was first described in the *M. incognita* genome (Mejias *et al.*, 2020; Nguyen *et al.*, 2018; Rutter *et al.*, 2014). Database queries showed that MiEFF18 displayed no sequence homology or known domains, and that it was absent from nematodes of other genera, such as cyst nematodes and free-living nematodes. By contrast, EFF18 orthologues were identified in seven of the eight RKNs for which genome sequences were available: *M. incognita*, *M. javanica*, *M. arenaria* (Blanc-Mathieu *et al.*, 2017), *M. hapla* (Opperman *et al.*, 2008), *M. enterolobii* (syn. *M. mayaguensis*) (Koutsovoulos *et al.*, 2020), *M. floridensis* (Lunt *et al.*, 2014) and *M. luci* (Susič *et al.*, 2020) (Figure 1, Table S1, Figure S1). No EFF18 orthologue was identified in *M. graminicola* (Somvanshi *et al.*, 2018). Three paralogous copies were identified, in the *M. incognita*, *M. javanica* and in *M. luci* genomes. Four copies were detected in *M. arenaria* and a single copy was detected in *M. hapla, M. floridensis* and *M. enterolobii*. A sequence alignment and analysis of the RKN EFF18 protein sequences showed that they were more than 60% identical, between 279 and 316 amino acids (aa) long and that they had an N-terminal secretion signal peptide (SSP), a low-complexity acidic D/E-rich region and a C-terminal lysine (K)-rich domain carrying direct repeats (Figure 1a, Figure S2). Only the C-terminal K-rich domain displayed marked differences between copies.

**Figure 1.**
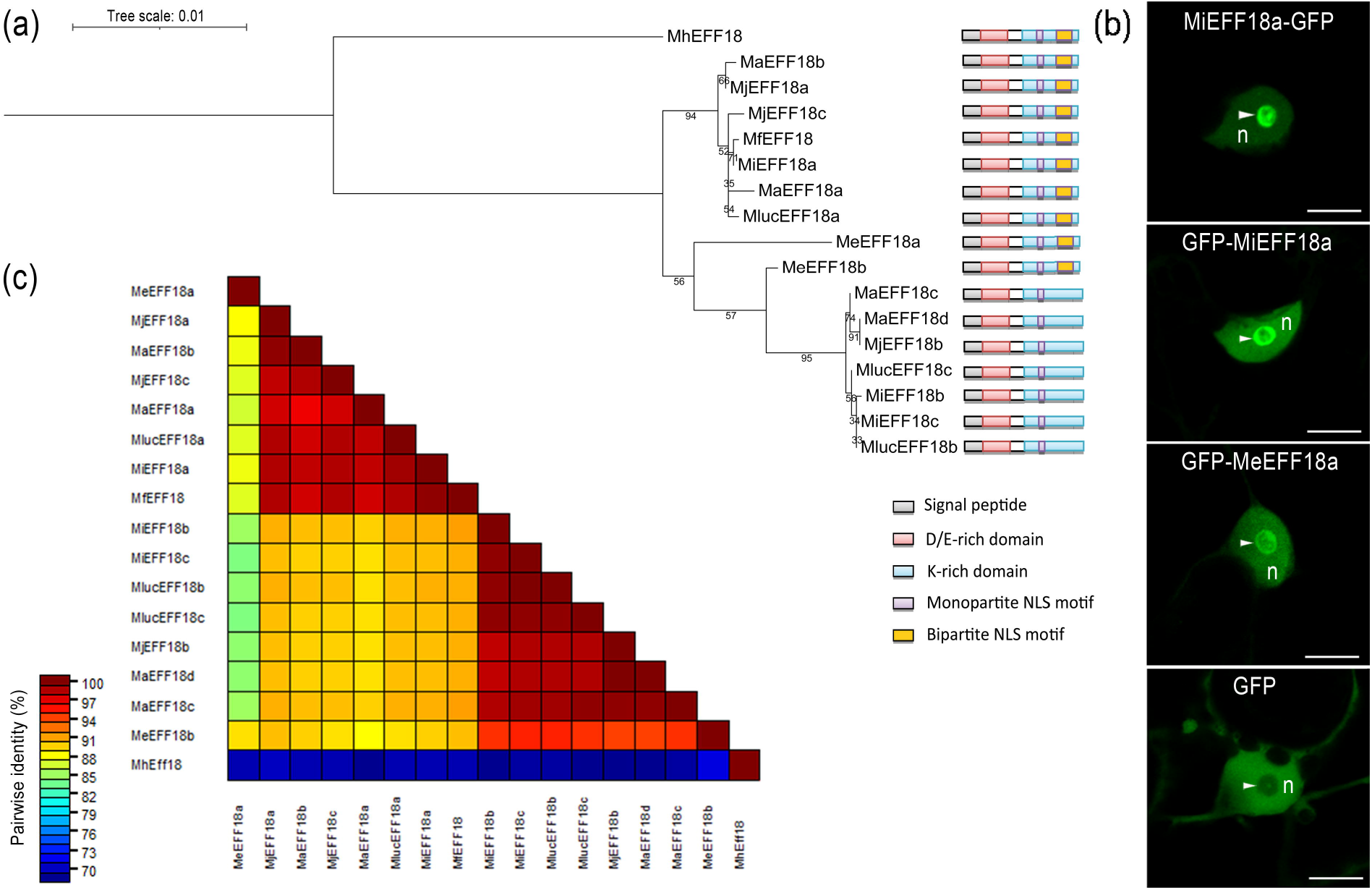
Effector 18 (EFF18) is a conserved effector in root-knot nematodes. (a) Phylogenetic tree and schematic diagram of root-knot nematode EFF18 protein sequences. The tree scale corresponds to the number of substitutions per site based on the amino-acid matrix (JTT). In the schematic diagram of EFF18 proteins, the predicted secretion signal peptide (SP; grey boxes), the aspartic acid and glutamic acid (D-E)-rich region (red boxes), the lysine (K)–rich C-terminal region (blue boxes) and the nuclear mono-(purple boxes) or bi-(orange boxes) partite localisation signals (NLS) are shown. EFF18 proteins from the closest group to MiEFF18a carry one mono- and one bipartite NLS, whereas the most divergent copies have only a single monopartite NLS. (b) MiEFF18 localised to the nucleus and nucleolus of plant cells. The MiEFF18 sequence was fused to that encoding GFP in an N- or C-terminal position and expressed in *N. benthamiana* leaves by agroinfiltration. GFP was used as a control and gave fluorescence in the cytoplasm and the nucleus (n), but not the nucleolus (arrowhead). Bars = 10 μm. (c) Pairwise sequence identity matrix for RKN EFF18 protein sequences.

A phylogenetic tree based on an alignment of the 17 RKN EFF18 protein sequences showed divergences between copies among the same species (Figure 1a). EFF18 proteins more closely related to MiEFF18a/Minc18636 harboured one monopartite NLS and one bipartite NLS, whereas other copies are more divergent (*e.g.* MiEFF18b/Minc15401 and MiEFF18c) and contained only one monopartite NLS (Figure 1a). MiEFF18a fused at its C- or N-terminus to GFP (green fluorescent protein) and MeEFF18a fused at its N-terminus to GFP were transiently expressed in *Nicotiana benthamiana* leaf epidermis. For EFF18 constructs, GFP fluorescence was only detected in the nucleus, with a strong GFP signal accumulating in the nucleolus (Figure 1b). In contrast, GFP alone was detected in the cytoplasm and the nucleus, but not in the nucleolus (Figure 1b). The EFF18s with bipartite NLS were 98% to 100% identical to the MiEFF18a protein, whereas those with only monopartite NLS were only 79 to 89% identical to this protein (Figure 1c). *M. hapla* had the most divergent genome of the *Meloidogyne* species tested. It was found to have a single copy of the gene, 63-65% identical to the closest copies and the most divergent copies, which suggests that the ancestor of RKN species had an EFF18 gene, and providing support for the role of EFF18 as a conserved effector.

### RKN EFF18s are specifically expressed in the subventral glands

MiEFF18 have been shown to be more strongly expressed at parasitic stages and to be expressed specifically in the subventral glands of *M. incognita* J2s (Rutter et al., 2014; Nguyen et al., 2018; Mejias et al., 2020). We studied the pattern of expression of genes encoding orthologous sequences of MiEFF18 in two other RKN species, by performing *in situ* hybridisation (ISH) for the *M. enterolobii MeEFF18a* and the *M. arenaria MaEFF18a* sequences. A specific signal was detected in the subventral oesophageal gland cells of pre-J2s after hybridisation with digoxigenin-labelled MeEFF18a and MaEFF18a antisense probes (Figure 2). No signal was detected in pre-J2s with sense negative controls. This finding suggests that MaEFF18a and MeEFF18a, may, like MiEFF18a, be secreted and play an important role in nematode parasitism.

**Figure 2.**
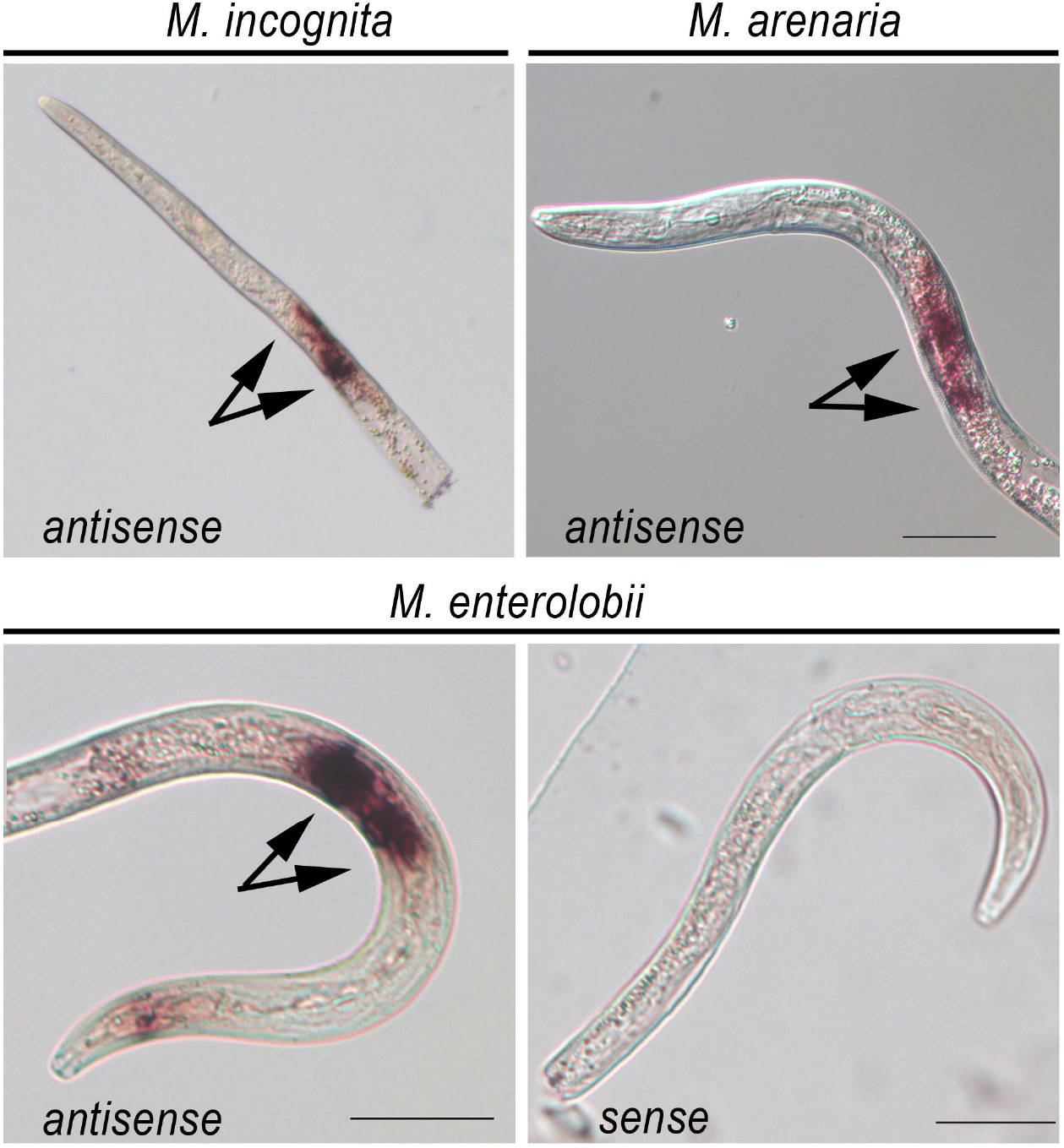
RKN EFF18s are specifically expressed in the subventral glands. *In situ* hybridisation, showing EFF18 transcripts in the subventral glands of J2s of *M. incognita*, *M. enterolobii* and *M. arenaria*, respectively. A sense probe for the MeEFF18 transcript was used as a negative control. SvG, subventral glands. Bar = 50 μm.

### MiEFF18a and MeEFF18a interact with the SmD1 proteins of *A. thaliana*, *N. benthamiana* and *S. lycopersicum*

We have demonstrated an interaction between MiEFF18 and the nuclear ribonucleoproteins SmD1s from *S. lycopersicum* and *A. thaliana,* modulating the pattern of alternative splicing and promoting the formation of giant cells (Mejias et al., 2020). Two genes, *AtSmD1a* (AT3G07590) and *AtSmD1b* (AT4G02840), encode SmD1 proteins in *Arabidopsis* (Koncz et al., 2012) and two genes encode 100% identical SmD1 proteins (SlSmD1) in *S. lycopersicum: SlSmD1a* (Solyc06g084310) and *SlSmD1b* (Solyc09g064660). In *N. benthamiana*, we identified three genes encoding SmD1s: *NbSmD1a* (*Niben101Scf01782g05006*), *NbSmD1b* (*Niben101Scf05290g01011*), and *NbSmD1c* (*Niben101Scf04283g03011*). A multiple sequence alignment showed that SmD1 was highly conserved in these species, with 93% identity between SlSmD1 and the sequence from which it diverged most strongly, AtSmD1b (Figure 3a). Like all Sm proteins, SmD1s carry two conserved Sm motifs mediating protein-protein interactions during small nuclear ribonucleoprotein (snRNP) biogenesis (Figure 3b). We investigated the subcellular localisation of SmD1 in plant cells, by transiently expressing constructs encoding GFP-SmD1 fusion proteins in *N. benthamiana*. We confirmed a strong accumulation of SlSmD1a and AtSmD1b in the nucleolus and in Cajal bodies, and a weaker accumulation in the nucleoplasm (Figure 3c).

**Figure 3.**
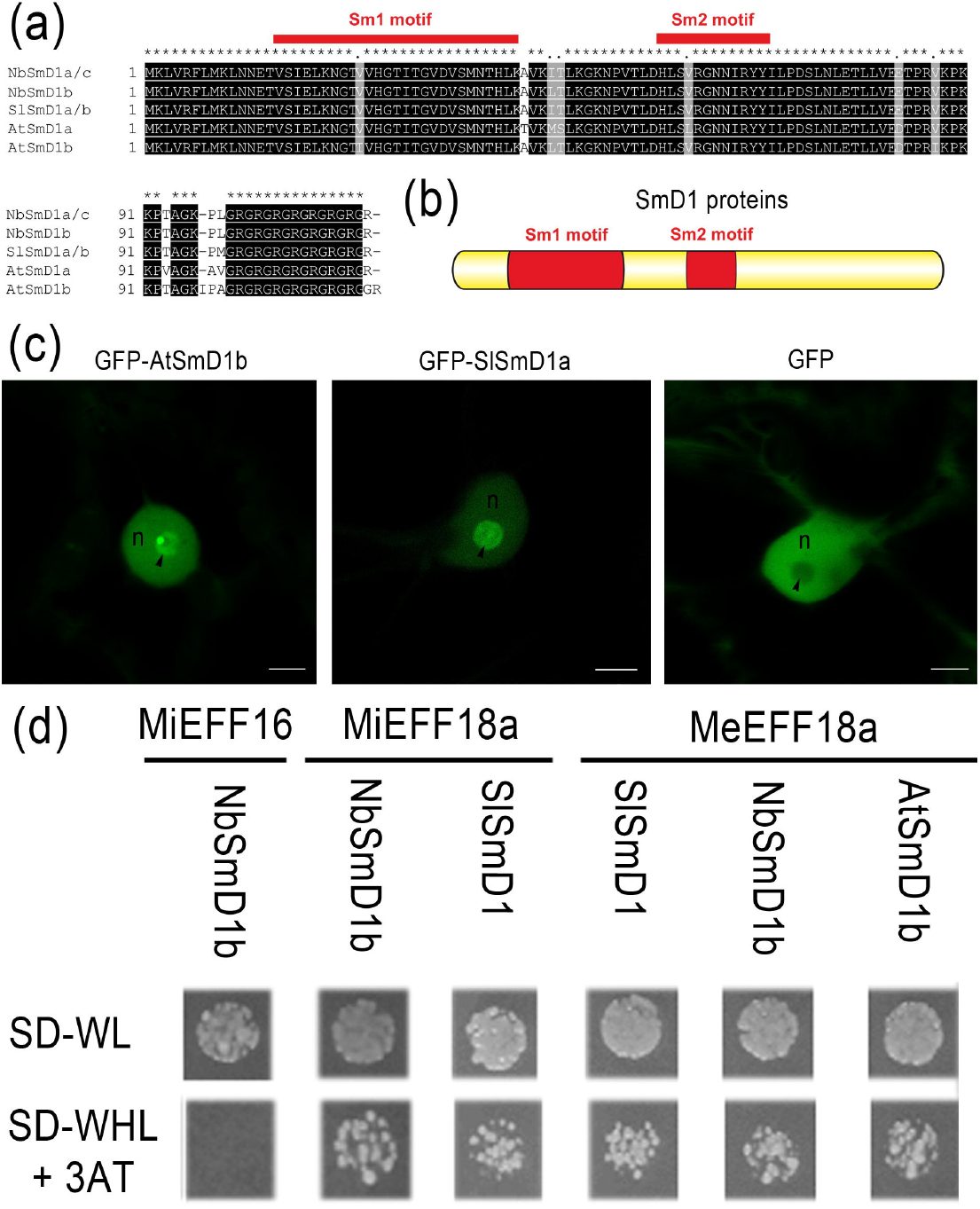
Conserved SmD1 proteins are targeted by EFF18. (a) MAFFT protein sequence alignment of the *S. lycopersicum* (Sl), *N. benthamiana* (Nb) and *A. thaliana* (At) SmD1 proteins. (b) Schematic representation of Sm1 and Sm2 motif in SmD1 proteins. (c) GFP-AtSmD1b and GFP-SlSmD1a accumulate in the nucleus and particularly in the nucleolus when transiently expressed in *N. benthamiana* epidermal leaf cells. GFP was used as a nucleocytoplasmic control. n= nucleoplasm; Black arrows show nucleolus. Bars = 5 μm. (d) Pairwise yeast two-hybrid assays showed that the MiEFF18 and MeEFF18 proteins were able to interact with the SmD1 proteins of *A. thaliana*, *S. lycopersicum* and *N. benthamiana*. We used MiEFF18 and MiEFF16 as a positive and negative control, respectively. Diploid yeasts containing the bait and prey plasmids carrying controls, effectors or SmD1 were serially diluted and spotted on plates. The 10-2 dilution is shown. SD-WL corresponds to the non-selective medium without tryptophan (W) and leucine (L). Only yeasts carrying a protein-protein interaction can survive on the SD-WLH (H, histidine) + 0.5 mM 3-aminotriazole (3-AT) selective medium.

We then investigated whether MeEFF18a was also able to interact with SmD1 proteins from *S. lycopersicum* and *A. thaliana,* like MiEFF18 (Mejias et al., 2020). Using a pairwise yeast-two hybrid approach, we showed that MiEFF18a and MeEFF18a interact with SmD1 proteins from plants of different clades, such as *A. thaliana*, *S. lycopersicum* and *N. benthamiana* (Figure 3d). As a control, we tested SmD1 interactions with another *M. incognita* effector, MiEFF16, encoded by the *Minc16401* gene and expressed in the subventral glands, with the same nuclear location *in planta* as MiEFF18 (Mejias et al., 2020). No interaction was observed between MiEFF16 and SmD1 proteins in yeast (Figure 3d). These results demonstrate that EFF18 proteins are conserved among RKNs and that they interact with SmD1 proteins, which are conserved among plant species.

### *SmD1* acts as a susceptibility gene for infection in plants of different clades

We recently demonstrated an important role for the AtSmD1b protein in giant cell formation and successful nematode infection (Mejias et al., 2020). We investigated whether *SmD1* is a conserved susceptibility gene required to ensure infection, and essential for RKN parasitism in Solanaceae species, by using a virus-induced gene silencing (VIGS) approach to alter the expression of *SmD1* genes in *S. lycopersicum* and *N. benthamiana*.

We first performed a VIGS assay to silence *SmD1* genes in *S. lycopersicum* (Figure 4a). We evaluated silencing efficiency by RT-qPCR on emerging leaves. Treated tomatoes had much lower levels of *SmD1* transcripts (Figure 4b). Tomatoes in which SmD1 genes were silenced displayed developmental defects on emerging leaves and had a shorter root system (Figure S3). In tomato plants infected with *M. incognita,* in which *SmD1* genes were silenced, the number of females producing egg masses was much smaller than that in control plants treated with the TRV-GFP virus (Figure 4c).

**Figure 4.**
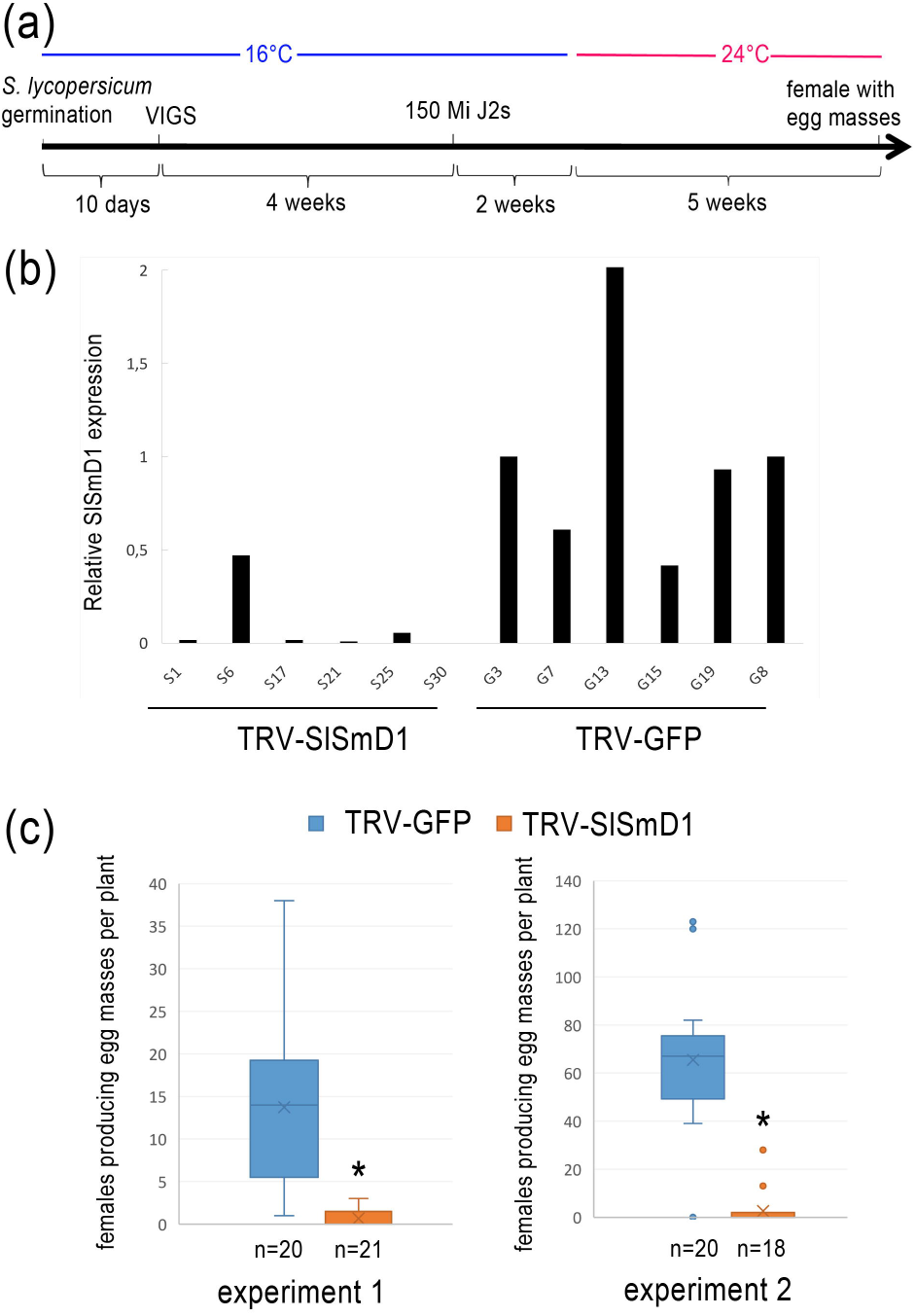
The silencing of SmD1 genes by VIGS affects susceptibility to *M. incognita* in *S. lycopersicum*. (a) Timeline used for the VIGS experiments in S. lycopersicum. (b) RT-qPCR demonstrating the effective silencing of *SlSmD1* in TRV-SlSmD1 line when compared to the control TRV-GFP. RPN7 was used for data normalisation. (c) Infection test on tomato plants in which SmD1 was silenced (TRV-SlSmD1) and control tomato plants (TRV-GFP). Females producing egg masses were counted seven weeks after inoculation with 150 *M. incognita* J2s per plant. Statistical significance was determined on the basis of three independent biological replicates, by Kruskal-Wallis tests. Clearly significant differences were observed between TRV-GFP control and TRV-SmD1 plants (*p ≤ 0.05).

Because of adverse effect of *SmD1* silencing on development in tomato, we then silenced the *SmD1* genes in *N. benthamiana,* which allows performing a VIGS assay at a later developmental stage when roots have already developed substantially (Figure 5a and 5b). An evaluation of silencing efficiency by RT-qPCR showed that *N. benthamiana* roots subjected to VIGS had much lower levels of *SmD1* transcripts, particularly for the most strongly expressed gene, *NbSmD1b* (Figure 5c). We observed no significant decrease in the expression of the two mostly weakly expressed genes, *NbSmD1a* and *NbSmD1c* (Figure 5c; Figure S4). *N. benthamiana* plants in which *SmD1* was silenced produced a much smaller number of galls (up to 80% fewer) following infection with *M. incognita* (Figure 5d).

**Figure 5.**
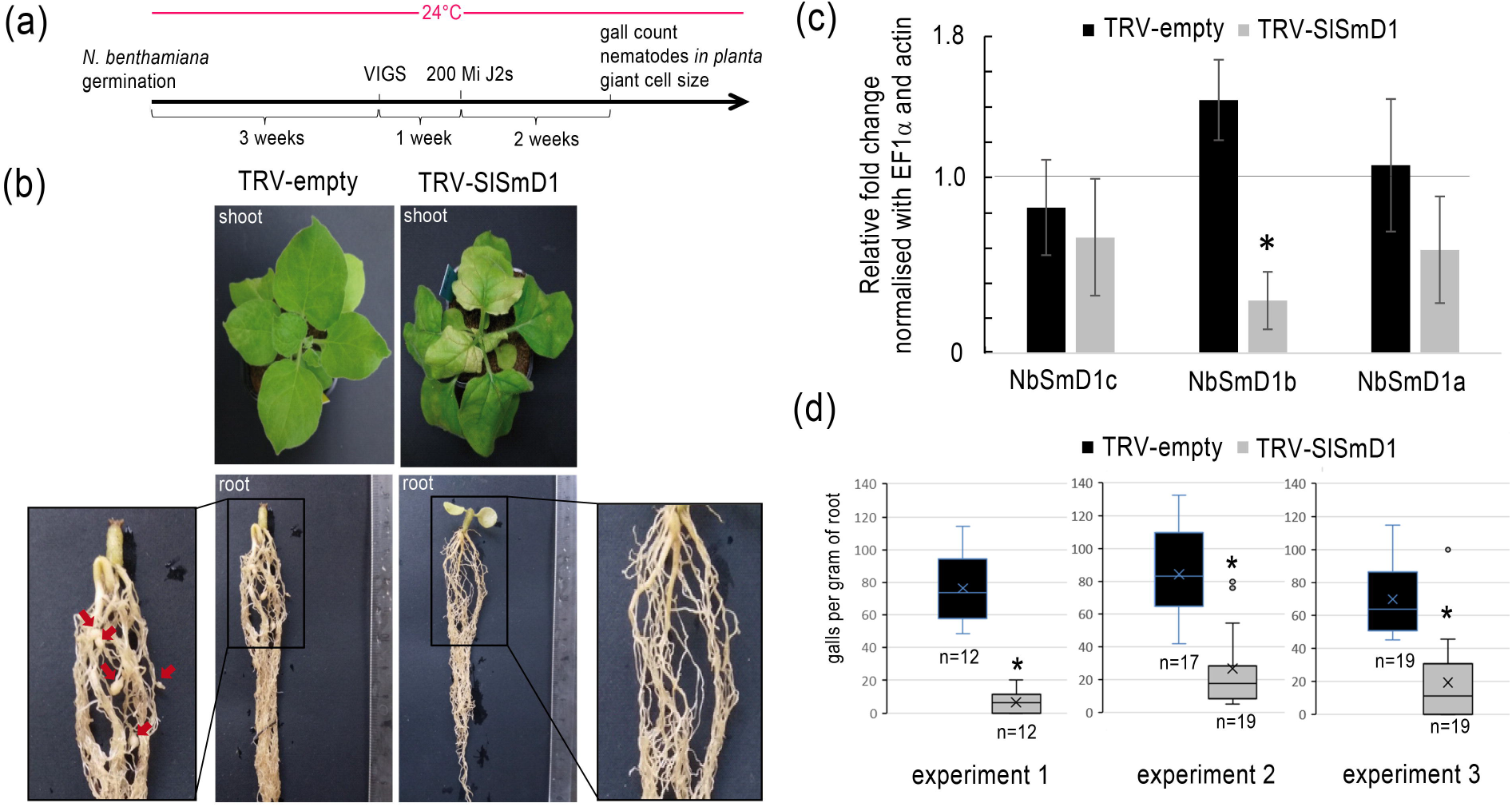
The silencing of *SmD1* genes by VIGS affects susceptibility to *M. incognita* in *N. benthamiana*. (a) Timeline used for VIGS experiment in *N. benthamiana*. (b) *N. benthamiana* plants with silenced SmD1 genes (TRV-SmD1, right panel) and TRV2-empty control plants (TRV-empty, left panel), showing some developmental defects of the leaves (upper panel) and a shorter root system (lower panel). Red arrow point-out galls on these pictures. (c) RT-qPCR showing that the NbSmD1b gene, the most strongly expressed and closest orthologue to the SlSmD1 gene, was effectively silenced. The data shown are the means of three independent biological replicates. EF1a and actin were used for data normalisation. (d) Infection test on *N. benthamiana* control plants (TRV-empty) and plants in which SmD1 was silenced (TRV-SlSmD1). Galls were counted and root weight was measured two weeks after inoculation with 200 *M. incognita* J2s per plant. Statistical significance was determined on the basis of three independent biological replicates, by Kruskal-Wallis tests. Clear significant differences were found between the TRV-GFP control and the TRV-SmD1 plants (*p ≤ 0.05; **p ≤ 0.01).

We studied the effect on nematode and giant cell development in detail, by investigating J2s *in planta* by the fuchsine acid staining method, to determine the proportions of migrating filiform and sedentary swollen parasitic juveniles and their ratio. The percentage of migrating filiform J2s was higher (90%) in plants in which *SmD1* was silenced, which had a lower percentage of swollen juveniles, indicating a defect in the RKN development (Figure 6a). We also investigated whether the giant cells formed on plants in which *SmD1* was silenced displayed developmental defects. We observed these cells directly, under a confocal microscope, after clearing in benzyl alcohol/benzyl benzoate (BABB; Cabrera et al., 2018). A comparison of the mean surface areas of the largest giant cells in each gall showed that giant cells from plants in which *SmD1* was silenced were 36% smaller than those from control plants (Figure 6b and 6c). These results confirm the important role of SmD1 in giant cell formation in Solanaceae species and the requirement of this protein for successful nematode development.

**Figure 6.**
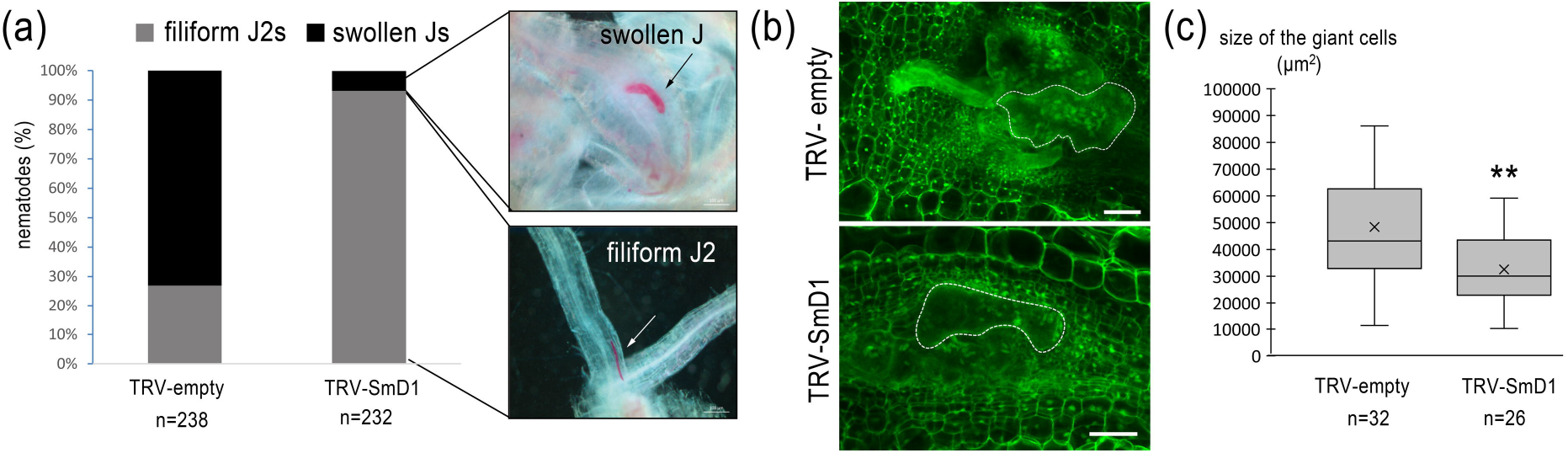
*SmD1* plays an important role in the formation of giant cells. (a) The filiform J2s/swollen juveniles (Js) ratio obtained by fuchsine acid staining in the *N. benthamiana* root system with (TRV-SmD1) and without (TRV-empty) silencing with the TRV-SlSmD1 construct, following infection with *M. incognita*. (b) Galls of negative control plants and plants with SmD1 silencing collected two weeks post infection for measurement of the area of giant cells (dotted line) by the BABB clearing method (Cabrera et al., 2018). The biggest giant cell measured is shown by a surrounding dashed white line. Bar = 100 μm. (c) Box-and-whisker plot of giant cell size (μm2) measurements (n = 32 and 26 galls).

## Discussion

The ability of plant pathogens to infect their hosts is generally dependent on the secretion of effectors. Most pathogens secrete effectors to overcome host physical defences, such as the plant cell wall, and to suppress plant immune responses (Toruño *et al.*, 2016). Other effectors are more specific to the parasitic strategy of the pathogen and may regulate host gene expression or trigger changes in host cell morphology and physiology to allow pathogen feeding and development. Most obligatory biotrophs form specific feeding structures, such as the haustoria of biotrophic filamentous pathogens, and produce sets of specific effectors (Chaudhari *et al.*, 2014; O’Connell and Panstruga, 2006). RKNs are root endoparasites that manipulate host cells to form specialised giant cells for feeding. These giant cells constitute the sole source of nutrients for the nematode, and are, therefore, essential for nematode survival. RKNs induce giant cells by manipulating root cell developmental programmes. Indeed, massive transcriptomic reprogramming occurs during giant cell formation (Favery *et al.*, 2016; Mitchum *et al.*, 2013). Genes associated with root meristem function, lateral root formation and the establishment of the vasculature, in particular, are tightly regulated upon giant cell induction (Cabrera *et al.*, 2014; Olmo *et al.*, 2020; Yamaguchi *et al.*, 2017). Alternative splicing has recently been shown to occur in *Arabidopsis* following infection with *M. incognita*, and this process contributes to transcriptome and proteome diversity (Mejias et al., 2020).

### EFF18 is a nuclear conserved RKN-specific effector

Nuclear effectors are thought to mediate the transcriptional reprogramming required for giant cell formation (Mejias *et al.*, 2019; Quentin *et al.*, 2013). They may interfere with the function of transcription factors, as described for Mi16D10, which interacts with SCARECROW-like transcription factors (Huang *et al.*, 2006), or may themselves act as transcription factors, as reported for Mi7H08 (Zhang *et al.*, 2015). MiEFF18 is another RKN effector that has been shown to be secreted within host cells, in which it localises to the nucleus. MiEFF18 has been shown to interact with the SmD1 protein, a core component of the spliceosome conserved in all eukaryotes, thereby modulating alternative splicing and gene expression (Mejias et al., 2020). MiEFF18 may corrupt the function of *Arabidopsis SmD1* function to modulate the expression of various genes encoding proteins involved in giant cell formation through processes such as DNA replication or cytokinesis (Mejias et al., 2020). We show here that the manipulation of SmD1 function by MiEFF18 plays a key role in giant cell development in other plant species, such as *Nicotiana benthamiana* and the tomato *Solanum lycopersicum.*

Genes encoding the MiEFF18 effector were found in all available *Meloidogyne* spp. genomes other than the draft genome for the rice RKN *M. graminicola* (Somvanshi *et al.*, 2018). EFF18 is exclusive to RKN, being absent from all other parasitic nematodes and other plant pathogens with parasitic strategies not involving the induction of giant feeding cells. At least one orthologous copy of a MiEFF18 sequence was detected in each of the available *Meloidogyne* genomes, demonstrating that MiEFF18 is a conserved effector. The multiple copies identified in *M. incognita, M. javanica* and *M. arenaria* are consistent with the polyploidy of these mitotic parthenogenetic species (Koutsovoulos *et al.*, 2020). The absence of an EFF18 effector in *M. graminicola* may be explained by the particular host range and life cycle of this nematode. *M. graminicola* may have lost the EFF18 effector during specialisation on monocotyledonous hosts and adaptation to the infection of submerged roots (Mantelin *et al.*, 2017). These growing conditions may have resulted in different requirements for the modulation of gene expression for giant cell ontogenesis, dependent on effectors other than EFF18.

The distribution of EFF18 orthologues in two major groups, with copies (e.g. MiEFF18a) carrying two NLS, and those of the most divergent group (e.g. MiEFF18b) carrying only one NLS, suggested a possible duplication of the ancestral MiEFF18 gene in the ancestor of RKN species, with one of the duplicated genes either gaining or losing a bipartite NLS. The proteins from the closest group to the MiEFF18a gene would be expected to function similarly to MiEFF18a, through the modulation of SmD1 functions, due to the very high level of sequence identity between these proteins (98% identity). *M. enterolobii* is an extremely polyphagous species that reproduces through mitotic parthenogenesis, like *M. incognita*. Therefore, we investigated the functionality of proteins MeEFF18 orthologue. We found that, like MiEFF18, MeEFF18a was able to interact with SmD1 proteins from *A. thaliana*, *N. benthamiana* and *S. lycopersicum*, suggesting that orthologous copies of MiEFF18a are functional and target the same functions in different host plants. MiEFF18a and MeEFF18a are the first examples of conserved RKN effectors able to target the same conserved plant process in different plant species.

### Targeting conserved effectors to engineer plant resistance

The identification of conserved effectors could lead to new strategies for developing broad resistance (Huang *et al.*, 2006; Landry *et al.*, 2020; Peeters *et al.*, 2013; Roux *et al.*, 2015). Only a few RKN effectors have been described to be conserved. The MAP (*Meloidogyne* avirulence protein) effector family, which includes *M. incognita* Mi-MAP1.2, was shown to be conserved in 13 of the 21 RKN species tested, and absent from other genera of plant-parasitic nematodes (PPNs) (Tomalova *et al.*, 2012). The genes of the MAP effector family harbour one or multiple CLE-like motifs, which may be involved in feeding site formation, as demonstrated for cyst nematode CLE-like peptides, which promote syncytium formation (Rutter et al., 2014; Mitchum et al., 2012). MjNULG1a, from *M. javanica,* is a nuclear effector with a demonstrated role in parasitism. Southern blot experiments have revealed the presence of MjNULG1a orthologues in *M. incognita* and *M. enterolobii*, but not in other PPNs (Lin *et al.*, 2013). Similarly, the 16D10 effector is exclusive to RKNs (Huang et al., 2006; Dinh, 2015). The use of host-induced gene silencing (HIGS) approaches to engineer plant resistance to RKNs has excited considerable interest (Ali *et al.*, 2017; Banerjee *et al.*, 2017). The targeting of genes involved in nematode development or encoding effectors has been considered. Silencing conserved effectors may allow specific resistance to RKNs with no impact on non-targeted species. Studies of Mi16D10 have demonstrated the feasibility of conferring RKN resistance in *Arabidopsis*, potato or grape through the targeting of this effector (Huang et al., 2006; Yang et al., 2013; Dinh, 2015). However, this strategy is constrained both by limited HIGS effectiveness, by the redundancy of the effector's function and the difficulty in targeting the point in time when the effector plays a key role in the interaction.

### Targeting essential conserved effector targets to induce a loss of susceptibility

The use of resistant cultivars or rootstocks is an efficient and non-polluting method for controlling RKNs. Very few natural resistance genes (R-genes) have been identified to date, in a limited number of plant species. Furthermore, some RKN species or populations are not controlled by these genes, e.g. *M. enterolobii* (Elling, 2013; Kiewnick *et al.*, 2009) or can overcome these resistances, e.g. populations of *M. incognita* virulent against tomato *Mi1.2* (Castagnone-Sereno, 2006). One alternative would be to target conserved plant genes encoding proteins involved in host processes that are hijacked by the biotrophic pathogens for settlement and feeding, and that are essential for disease development. These susceptibility genes (S-genes) represent an alternative to R-genes for the deployment of pathogen resistance, and they may be more durable in the field (Dong and Ronald, 2019; Engelhardt *et al.*, 2018; van Schie and Takken, 2014). Well-characterised examples of S-genes include the genes encoding eukaryotic translation initiation factors, the sugar transporter SWEET14 or PMR6, which are required for viral, bacterial and oomycete infections, respectively (Langner *et al.*, 2018; van Schie and Takken, 2014; Schmitt-Keichinger, 2019).

In recent decades, transcriptomic approaches have been widely used to identify genes regulated upon RKN infection, and, thus, host functions manipulated by RKNs. However, as thousands of genes are differentially regulated during a compatible interaction, the identification of S-genes from these data is a very time-consuming process, probably explaining why only a few genes to date have been shown to be important for the establishment of giant cells (Favery *et al.*, 2016). Interactomics approaches have recently been used to identify the direct plant targets manipulated by PPN effectors. Only a few targets of RKN effectors have been identified, but most have been shown to be instrumental in promoting nematode parasitism (Mejias *et al.*, 2019). SmD1 is a host target of an effector required for host susceptibility to RKNs in several plant clades. It exerts a conserved plant function targeted by a core effector in *Arabidopsis* and Solanaceae, common to diverse *Meloidogyne* species that have adopted the same successful parasitic strategy based on the induction of giant feeding cells in the root in several host species. SmD1 is thus a good candidate S-gene for targeting in the engineering of crop resistance to RKN. As SmD1 functions is required for plant development knockout mutations of this gene have adverse effects, it will be necessary to identify mutant alleles that can evade recognition by MiEFF18 whilst remaining competent to perform the functions of SmD1 in the regulation of plant development in a continually changing environment. This strategy has proven to be effective for potyvirus susceptibility *eIF4E* genes (Bastet *et al.*, 2019).

## Experimental procedures

### Nematode and plant materials

*M. incognita* (Morelos strain), *M. arenaria* (Guadeloupe strain) and *M. enterolobii* (Godet strain) were multiplied on tomato (*Solanum lycopersicum* cv. “Saint Pierre”) growing in a growth chamber (25°C, 16 h photoperiod). Freshly hatched J2s were collected as previously described (Caillaud and Favery, 2016). For VIGS experiments, *Nicotiana benthamiana* and *Solanum lycopersicum* (cv M82) seeds were sown on soil and incubated at 4°C for two days. After germination, *N. benthamiana* and tomato plantlets were transplanted into pots containing soil and sand (1:1), and were grown at 24°C and 16°C, respectively (photoperiod, 16 h: 8 h, light: dark).

### Sequence analysis, alignment and phylogenetic tree

The sequences of EFF18 paralogues and orthologues were obtained from *Meloidogyne* genomic resources http://www6.inra.fr/meloidogyne_incognita and Wormbase parasite. Protein sequences were aligned with the MAFFT tool on the EBI server (https://www.ebi.ac.uk/Tools/msa/mafft/). The alignment was then used as input for the IQTree Web server http://iqtree.cibiv.univie.ac.at/ (Trifinopoulos et al., 2016) to generate the maximum likelihood phylogenetic tree. The model chosen by the inbuilt model test was HIVbr+F+G4. Support for the nodes was calculated with 100 bootstrap replicates. *M. hapla* was used as the outgroup in the phylogenetic tree for MiEFF18 orthologues. The tree was visualised in iTOL https://itol.embl.de/. The sequence alignment were coloured with Boxshade (https://embnet.vital-it.ch/software/BOX_form.html). The pairwise sequence identity matrix of RKN EFF18 protein sequences was generated with Sequence Demarcation Tool version 1.2 software (Muhire *et al.*, 2014) (http://web.cbio.uct.ac.za/~brejnev/).

### *In situ* hybridisation (ISH)

ISH was performed on freshly hatched *M. arenaria* and *M. enterolobii* pre-J2s, as previously described (Jaouannet *et al.*, 2018). The MaEFF18, and MeEFF18 gene fragments were amplified with specific primers (Table S2). A sense probe for MeEFF18 was used as a negative control. Images were obtained with a microscope (Zeiss Axioplan2, Germany).

### Pairwise yeast two-hybrid assays

For pairwise yeast two-hybrid (Y2H) assays, the coding sequences (CDS) of the MiEFF16, MiEFF18 and MeEFF18 effectors without their secretion signals were amplified (Table S1) and inserted into pB27 as C-terminal fusions with LexA. Full-length SmD1 CDS sequences (*SlSmD1*, *NbSmD1* and AtSmD1b) were amplified (Table S1) and inserted into pP6 as C-terminal fusions with Gal4-AD. The pB27 and pP6 constructs were verified by sequencing and used to transform the L40ΔGal4 (MATa) and Y187 (MATα) yeast strains, respectively. Y187 and L40ΔGal4 were crossed and diploids were selected on medium lacking tryptophan and leucine. Interactions were investigated on medium lacking tryptophan, leucine and histidine and supplemented with 0.5 mM 3-aminotriazole (3-AT).

### *N. benthamiana* agroinfiltration

Transient expression was achieved by infiltrating *N. benthamiana* leaves with *A. tumefaciens* GV3101 strains harbouring GFP-fusions, as previously described (Caillaud *et al.*, 2008). Leaves were imaged 48 hours after agroinfiltration, with an inverted confocal microscope equipped with an argon ion and HeNe laser as the excitation source. For simultaneous GFP imaging, samples were excited at 488 nm and GFP emission was detected selectively with a 505-530 nm band-pass emission filter.

### Virus-induced gene silencing in Solanaceae

VIGS assays were performed on *N. benthamiana* and *S. lycopersicum*. We used the Sol Genomics Network VIGS-Tool (https://vigs.solgenomics.net/) to design the best sequence for silencing SlSmD1a transcripts, and selected the full-length *SlSmD1a* (without the ATG and STOP codons) for amplification by PCR with the TRV2-SlSmD1-F/TRV2-SlSmD1-R primer pairs (Table S2). The PCR products were digested with *Eco*RI and *Xho*I and ligated to the tobacco rattle virus RNA 2 vector (TRV2) for the transformation of *A. tumefaciens* strain GV3101. VIGS assays were performed, as previously described, by the co-infiltration of leaves of three-week-old *N. benthamiana* plants (Lange *et al.*, 2013; Velasquez *et al.*, 2009) or 10-days-old tomato plants (Cox *et al.*, 2019) with agrobacterial strains containing the RNA 1 vector (TRV1) and TRV2. Tomato plants were incubated at 16°C for four weeks. Three independent biological replicates were established for each set of conditions (*n* = 15 per replicate). Two *N. benthamiana* root systems per set of conditions and per replicate, or *S. lycopersicum* leaves were collected upon RKN infection and frozen in liquid nitrogen for subsequent RNA extraction and the assessment of silencing efficiency by RT-qPCR.

### RKN infection assay, juveniles in the plant and giant cell area measurements

*N. benthamiana* plants subjected to VIGS were inoculated with 200 *M. incognita* J2s per plant, 10 days post inoculation (dpi) with TRV, and incubated at 24°C. *S. lycopersicum* plants subjected to VIGS were inoculated with 150 *M. incognita* J2s per plant, 30 dpi with TRV, and incubated at 16°C for two weeks before transfer to 24°C. *N. benthamiana* infected roots were collected two weeks after infection whereas *S. lycopersicum* infected roots were collected six weeks after infection. Galls, and egg masses stained with 4.5% eosin were counted under a binocular microscope, and root system was weighted (*n*=12 to 19 and n=18 to 21 plants per replicates for *N. benthamiana* and *S. lycopersicum*, respectively). Three and two independent biological replicates were established for each set of conditions in *N. benthamiana* and *S. lycopersicum*, respectively. The impact of the plant lines on the number of galls per mg of root in *N. benthamiana* and the number of egg masses per plant in *S. lycopersicum* were analyzed using Kruskal Wallis test since the dependent variable did not follow a Normal distribution using a Shapiro-Wilk Test. The different replicates of the numbers of galls per mg of roots in *N. benthamiana* were pooled for the analyzes because there was no difference between the replications (X^2^_2_= 2.8, *P* = 0.248). By contrast, the different replicates of the number of egg masses per plant in *S. lycopersicum* varied depending on the replication (X^2^_1_= 5.3, *P* = 0.022), and they were analyzed separately. Thus, both the number of galls per mg of root in *N. benthamiana* and the number of egg masses per plant in *S. lycopersicum* varied significantly between the two plant lines tested (X^2^_1_= 57.2, *P* < 0.001; X^2^_1_ > 25.6, *P* < 0.001, respectively). For determination of the ratio of filiform-to-swollen nematodes, infected roots were collected 14 dpi, parasitic nematodes were stained with fuchsine acid, as previously described (Karssen and Moens, 1983), and nematodes were examined under a binocular microscope (model LSM 880; Zeiss) (*n*= 3 plants per replicate for TRV-empty lines and n = 5 plants per replicate for TRV-SmD1 lines, with a mean of 75 nematodes observed per condition and per replicate). Three independent biological replicates were established for each set of conditions. Statistical analyses were carried out with R software (R Development Core Team, version 3.1.3). For giant cell area measurements, galls were cleared in BABB, as previously described (Cabrera *et al.*, 2018), and examined under an inverted confocal microscope (model LSM 880; Zeiss). The mean areas of the biggest giant cell in each gall from *N. benthamiana*, for each genotype, were measured with Zeiss ZEN software (*n* = 32 and 26 galls for control and SmD1-VIGSed plants, respectively). One biological experiment was performed for giant cells measurement. The data were analysed with a *t*-test since the data followed a normal distribution (t = 3.5, *P* < 0.001).

### Reverse transcription-quantitative PCR

We assessed silencing efficiency in Solanaceae species, by extracting total RNA with TriZol (Invitrogen) and subjecting 1 μg of total RNA to reverse transcription with the Superscript IV reverse transcriptase (Invitrogen). qPCR analyses were performed as described by Nguyen et al., 2016. Primers were designed to amplify both SlSmD1 transcripts or to discriminate between the three copies of SmD1 present in *N. benthamiana* specifically according to their UTR, to prevent the amplification of TRV2-SlSmD1 constructs (Table S2). We performed RT-qPCR in triplicate on cDNA samples from three independent biological replicates. The *EF-1*α and Actin genes were used for the normalisation of RT-qPCR data (Liu *et al.*, 2012). Quantifications and statistical analyses were performed with SATqPCR (Rancurel *et al.*, 2019), and the results are expressed as normalised absolute quantities.

### Graphs and statistical analysis

Graphs and plots were created with Microsoft® Office Excel® 2016 and statistical calculations were performed in R.

### Accession numbers

The sequence data from this article can be found in the *Arabidopsis* Information Resource (https://www.arabidopsis.org/), Solgenomics (https://solgenomics.net/) and/or GenBank/EMBL databases. All RKN EFF18 protein sequences are presented in the Figure S1 The accession numbers are summarised in Table S1 including MiEFF18a (KX907770), MeEFF18a (MW272456), NbSmD1a (MT683762), NbSmD1b (MT683763) and NbSmD1c (MT683764).

## Acknowledgements

We thank Dr Johnathan Dalzell and Steven Dyer (Queen's University Belfast, UK) for tomato VIGS protocol and vectors, Pr S.P. Dinesh-Kumar (University of California, Davis, USA) for VIGS vectors and Hybrigenics Services (Paris, France) for providing the pB27 and pP6 vectors and the L40ΔGal4 and Y187 yeast strains. We thank Dr Nemo Peeters and Dr Laurent Deslandes (LIPM, Castanet Tolosan, France) and Gregori Bonnet (Syngenta seeds) for helpful discussions. Microscopy work was performed at the SPIBOC imaging facility of Institut Sophia Agrobiotech. We thank Dr Lucie S. Monticelli for providing the statistical analyses. We thank Dr Olivier Pierre and the whole platform team for their help with microscopy. This work was funded by the INRAE SPE department and the French Government (National Research Agency, ANR) through the ‘Investments for the Future’ LabEx SIGNALIFE: programme reference #ANR-11-LABX-0028-01, by the INRA-Syngenta Targetome project, by the French-Japanese bilateral collaboration programmes PHC SAKURA 2016 #35891VD and 2019 #43006VJ and by the French-Chinese bilateral collaboration program PHC XU GUANGQI 2020 #45478PF. J.M. holds a doctoral fellowship from the French *Ministère de l*◻*Enseignement Supérieur, de la Recherche et de l*◻*Innovation* (MENRT grant). N.M.T. was supported by a USTH fellowship, as part of the 911-USTH programme of the Ministry of Education and Training of The Socialist Republic of Vietnam. Y.P.C. obtained scholarships from the China Scholarship Council (No. 201806350108) for studies at INRAE, France.

## Author contributions

J.M. designed and performed experiments, and interpreted data; J.M., YC and N.M.T. generated constructs and performed yeast two-hybrid analysis; YC performed *in situ* hybridisation and *in planta* subcellular localisation experiments; KM performed VIGS in tomato and nematode infection tests; J.M., P.A., S.J.P., B.F. and M.Q. wrote the manuscript; B.F. and M.Q. obtained funding, designed the work and supervised the experiments and data analyses; all the authors read and edited the manuscript.

## Supporting information

**Figure S1** Identified EFF18 sequences in RKN species.

**Figure S2** EFF18 is a conserved RKN effector.

**Figure S3** Tomato phenotypes associated with VIGS of *SlSmD1* genes.

**Figure S4** Semi-quantitative RT-PCR expression analysis of *NbSmD1s* in *N. benthamiana* roots VIGS experiments.

**Table S1** Sequences used in this study and accession numbers

**Table S2** Primers used in this study

## References

Ali, M.A., Azeem, F., Abbas, A., Joyia, F.A., Li, H., and Dababat, A.A. (2017) Transgenic Strategies for Enhancement of Nematode Resistance in Plants. Frontiers in Plant Science, 8, 1–13.

de Almeida Engler, J. and Gheysen, G. (2013) Nematode-induced endoreduplication in plant host cells: why and how? Molecular Plant-Microbe Interactions, 26, 17–24.

Banerjee, S., Banerjee, A., Gill, S.S., Gupta, O.P., Dahuja, A., Jain, P.K., and Sirohi, A. (2017) RNA Interference: A Novel Source of Resistance to Combat Plant Parasitic Nematodes. Frontiers in Plant Science, 8:834, doi: 10.3389/fpls.2017.00834.

Bastet, A., Zafirov, D., Giovinazzo, N., Guyon-Debast, A., Nogué, F., Robaglia, C., and Gallois, J.L. (2019) Mimicking natural polymorphism in eIF4E by CRISPR-Cas9 base editing is associated with resistance to potyviruses. Plant Biotechnology Journal, 17, 1736–1750.

Blanc-Mathieu, R., Perfus-Barbeoch, L., Aury, J.-M.M., Da Rocha, M., Gouzy, J., Sallet, E., et al. (2017) Hybridization and polyploidy enable genomic plasticity without sex in the most devastating plant-parasitic nematodes. PLoS Genetics, 13, e1006777.

Blok, V.C., Jones, J.T., Phillips, M.S., and Trudgill, D.L. (2008) Parasitism genes and host range disparities in biotrophic nematodes: The conundrum of polyphagy versus specialisation. Bioessays, 30, 249–259.

Cabrera, J., Bustos, R., Favery, B., Fenoll, C., and Escobar, C. (2014) NEMATIC: a simple and versatile tool for the in silico analysis of plant-nematode interactions. Molecular Plant Pathology, 15, 627–636.

Cabrera, J., Olmo, R., Ruiz-Ferrer, V., Abreu, I., Hermans, C., Martinez-Argudo, I., et al. (2018) A Phenotyping Method of Giant Cells from Root-Knot Nematode Feeding Sites by Confocal Microscopy Highlights a Role for CHITINASE-LIKE 1 in Arabidopsis. International Journal of Molecular Sciences, 19, 429.

Caillaud, M.-C., Abad, P., and Favery, B. (2008) Cytoskeleton reorganization. Plant Signaling & Behavior, 3, 816–818.

Caillaud, M.-C.C. and Favery, B. (2016) In Vivo Imaging of Microtubule Organization in Dividing Giant Cell. In: Methods in Molecular Biology (Caillaud, M.-C., ed), pp. 137–144. New York: Springer New York.

Castagnone-Sereno, P. (2006) Genetic variability and adaptive evolution in parthenogenetic root-knot nematodes. Heredity, 96, 282–289.

Chaudhari, P., Ahmed, B., Joly, D.L., and Germain, H. (2014) Effector biology during biotrophic invasion of plant cells. Virulence, 5, 703–709.

Cox, D.E., Dyer, S., Weir, R., Cheseto, X., Sturrock, M., Coyne, D., et al. (2019) ABC transporter genes ABC-C6 and ABC-G33 alter plant-microbe-parasite interactions in the rhizosphere. Scientific Reports, 9, 19899.

Danchin, E.G.J., Rosso, M.-N.N., Vieira, P., de Almeida-Engler, J., Coutinho, P.M., Henrissat, B., and Abad, P. (2010) Multiple lateral gene transfers and duplications have promoted plant parasitism ability in nematodes. Proc Natl Acad Sci U S A, 107, 17651–17656.

Dong, O.X. and Ronald, P.C. (2019) Genetic Engineering for Disease Resistance in Plants: Recent Progress and Future Perspectives. Plant Physiology, 180, 26–38.

Doyle, E.A. and Lambert, K.N. (2003) Meloidogyne javanica chorismate mutase 1 alters plant cell development. Mol Plant Microbe Interact, 16, 123–131.

Elling, A. a (2013) Major emerging problems with minor meloidogyne species. Phytopathology, 103, 1092–1102.

Engelhardt, S., Stam, R., and Hückelhoven, R. (2018) Good Riddance? Breaking Disease Susceptibility in the Era of New Breeding Technologies. Agronomy, 8, 114.

Escobar, C., Barcala, M., Cabrera, J., and Fenoll, C. (2015) Overview of Root-Knot Nematodes and Giant Cells. In: Advances in Botanical Research, pp. 1–32.

Favery, B., Quentin, M., Jaubert-Possamai, S., and Abad, P. (2016) Gall-forming root-knot nematodes hijack key plant cellular functions to induce multinucleate and hypertrophied feeding cells. Journal of Insect Physiology, 84, 60–69.

Gleason, C., Polzin, F., Habash, S.S., Zhang, L., Utermark, J., Grundler, F.M.W., and Elashry, A. (2017) Identification of two Meloidogyne hapla genes and an investigation of their roles in the plant-nematode interaction. Molecular Plant-Microbe Interactions, 30, 101–112

Hewezi, T. and Baum, T.J. (2013) Manipulation of Plant Cells by Cyst and Root-Knot Nematode Effectors. Molecular Plant-Microbe Interactions, 26, 9–16.

Holbein, J., Franke, R.B., Marhavý, P., Fujita, S., Górecka, M., Sobczak, M., et al. (2019) Root endodermal barrier system contributes to defence against plant-parasitic cyst and root-knot nematodes. The Plant Journal, 100, 221–236.

Huang, G.Z., Allen, R., Davis, E.L., Baum, T.J., and Hussey, R.S. (2006) Engineering broad root-knot resistance in transgenic plants by RNAi silencing of a conserved and essential root-knot nematode parasitism gene. Proceedings of the National Academy of Sciences of the United States of America, 103, 14302–14306.

Jaouannet, M., Nguyen, C.-N., Quentin, M., Jaubert-Possamai, S., Rosso, M.-N., and Favery, B. (2018) In situ Hybridization (ISH) in Preparasitic and Parasitic Stages of the Plant-parasitic Nematode Meloidogyne spp. Bio-Protocol, 8, 1–13.

Jones, J.T., Haegeman, A., Danchin, E.G.J.J., Gaur, H.S., Helder, J., Jones, M.G.K.K., et al. (2013) Top 10 plant-parasitic nematodes in molecular plant pathology. Molecular Plant Pathology, 14, 946–961.

Karssen, G. and Moens, M. (1983) Root-knot nematodes. In: Plant nematology, pp. 59–90. Wallingford: CABI.

Kiewnick, S., Dessimoz, M., and Franck, L. (2009) Effects of the Mi-1 and the N root-knot nematode-resistance gene on infection and reproduction of Meloidogyne enterolobii on tomato and pepper cultivars. Journal of nematology, 41, 134–9.

Koutsovoulos, G.D., Poullet, M., Elashry, A., Kozlowski, D.K.L., Sallet, E., Da Rocha, M., et al. (2020) Genome assembly and annotation of Meloidogyne enterolobii, an emerging parthenogenetic root-knot nematode. Scientific Data, 7, 324.

Landry, D., González-Fuente, M., Deslandes, L., and Peeters, N. (2020) The large, diverse, and robust arsenal of Ralstonia solanacearum type III effectors and their in planta functions. Molecular Plant Pathology, 21, 1377–1388.

Lange, M., Yellina, A.L., Orashakova, S., and Becker, A. (2013) Virus-Induced Gene Silencing (VIGS) in Plants: An Overview of Target Species and the Virus-Derived Vector Systems. In: Methods in Molecular Biology, pp. 1–14. Humana Press Inc.

Langner, T., Kamoun, S., and Belhaj, K. (2018) CRISPR Crops: Plant Genome Editing Toward Disease Resistance. Annual Review of Phytopathology, 56, 479–512.

Lin, B., Zhuo, K., Chen, S., Hu, L., Sun, L., Wang, X., et al. (2016) A novel nematode effector suppresses plant immunity by activating host reactive oxygen species-scavenging system. New Phytologist, 209, 1159–1173.

Lin, B., Zhuo, K., Wu, P., Cui, R., Zhang, L.-H., and Liao, J. (2013) A novel effector protein, MJ-NULG1a, targeted to giant cell nuclei plays a role in Meloidogyne javanica parasitism. Molecular plant-microbe interactions, 26, 55–66.

Liu, D., Shi, L., Han, C., Yu, J., Li, D., and Zhang, Y. (2012) Validation of Reference Genes for Gene Expression Studies in Virus-Infected Nicotiana benthamiana Using Quantitative Real-Time PCR. PLoS ONE, 7, e46451.

Lunt, D.H., Kumar, S., Koutsovoulos, G., and Blaxter, M.L. (2014) The complex hybrid origins of the root knot nematodes revealed through comparative genomics. PeerJ, 2, e356.

Mantelin, S., Bellafiore, S., and Kyndt, T. (2017) Meloidogyne graminicolaL: a major threat to rice agriculture. Molecular Plant Pathology, 18, 3–15.

Mejias, J., Bazin, J., Truong, N., Chen, Y., Marteu, N., Bouteiller, N., et al. (2020) The root-knot nematode effector MiEFF18 interacts with the plant core spliceosomal protein SmD1 required for giant cell formation. New Phytologist, in press, doi: 10.1111/nph.17089.

Mejias, J., Truong, N.M., Abad, P., Favery, B., and Quentin, M. (2019) Plant Proteins and Processes Targeted by Parasitic Nematode Effectors. Frontiers in Plant Science, 10, 970.

Mitchum, M.G., Hussey, R.S., Baum, T.J., Wang, X., Elling, A.A., Wubben, M., and Davis, E.L. (2013) Nematode effector proteins: an emerging paradigm of parasitism. New Phytologist, 199, 879–894.

Mitchum, M.G., Wang, X., Wang, J., and Davis, E.L. (2012) Role of Nematode Peptides and Other Small Molecules in Plant Parasitism. Annual Review of Phytopathology, 50, 175–195.

Muhire, B.M., Varsani, A., and Martin, D.P. (2014) SDT: A Virus Classification Tool Based on Pairwise Sequence Alignment and Identity Calculation. PLoS ONE, 9, e108277.

Nguyen, C.-N., Perfus-Barbeoch, L., Quentin, M., Zhao, J., Magliano, M., Marteu, N., et al. (2018) A root-knot nematode small glycine and cysteine-rich secreted effector, MiSGCR1, is involved in plant parasitism. New Phytologist, 217, 687–699.

O’Connell, R.J. and Panstruga, R. (2006) Tete a tete inside a plant cell: establishing compatibility between plants and biotrophic fungi and oomycetes. New Phytologist, 171, 699–718.

Olmo, R., Cabrera, J., Díaz-Manzano, F.E., Ruiz-Ferrer, V., Barcala, M., Ishida, T., et al. (2020) Root-knot nematodes induce gall formation by recruiting developmental pathways of post-embryonic organogenesis and regeneration to promote transient pluripotency. New Phytologist, 227, 200–215.

Opperman, C.H., Bird, D.M., Williamson, V.M., Rokhsar, D.S., Burke, M., Cohn, J., et al. (2008) Sequence and genetic map of Meloidogyne hapla: A compact nematode genome for plant parasitism. Proc Natl Acad Sci U S A, 105, 14802–14807.

P. T. Y. Dinh (2015) Plant-nematode interactions and the application of RNA interference for controlling root-knot nematodes. PhD, 3, 54–67.

Peeters, N., Carrère, S., Anisimova, M., Plener, L., Cazalé, A.-C., and Genin, S. (2013) Repertoire, unified nomenclature and evolution of the Type III effector gene set in the Ralstonia solanacearum species complex. BMC Genomics, 14, 859.

Quentin, M., Abad, P., and Favery, B. (2013) Plant parasitic nematode effectors target host defense and nuclear functions to establish feeding cells. Frontiers in Plant Science, 4, 53.

Rancurel, C., van Tran, T., Elie, C., and Hilliou, F. (2019) SATQPCR: Website for statistical analysis of real-time quantitative PCR data. Molecular and Cellular Probes, 46, 101418.

Rodiuc, N., Vieira, P., Banora, M.Y., and de Almeida Engler, J. (2014) On the track of transfer cell formation by specialized plant-parasitic nematodes. Frontiers in Plant Science, 5, 1–14.

Roux, B., Bolot, S., Guy, E., Denancé, N., Lautier, M., Jardinaud, M.-F., et al. (2015) Genomics and transcriptomics of Xanthomonas campestris species challenge the concept of core type III effectome. BMC Genomics, 16, 975.

Rutter, W.B., Hewezi, T., Abubucker, S., Maier, T.R., Huang, G., Mitreva, M., et al. (2014) Mining Novel Effector Proteins from the Esophageal Gland Cells of Meloidogyne incognita. Molecular plant-microbe interactions, 27, 965–974.

van Schie, C.C.N. and Takken, F.L.W. (2014) Susceptibility Genes 101: How to Be a Good Host. Annual Review of Phytopathology, 52, 551–581.

Schmitt-Keichinger, C. (2019) Manipulating Cellular Factors to Combat Viruses: A Case Study From the Plant Eukaryotic Translation Initiation Factors eIF4. Frontiers in Microbiology, 10:17. doi: 10.3389/fmicb.2019.00017

Singh, S.K., Hodda, M., and Ash, G.J. (2013) Plant-parasitic nematodes of potential phytosanitary importance, their main hosts and reported yield losses. EPPO Bulletin, 43, 334–374.

Somvanshi, V.S., Tathode, M., Shukla, R.N., and Rao, U. (2018) Nematode Genome Announcement: A Draft Genome for Rice Root-Knot Nematode, Meloidogyne graminicola. Journal of Nematology, 50, 111–116.

Susič, N., Koutsovoulos, G.D., Riccio, C., Danchin, E.G.J., Blaxter, M.L., Lunt, D.H., et al. (2020) Genome sequence of the root-knot nematode Meloidogyne luci. Journal of Nematology, 52, e2020–25.

Tomalova, I., Iachia, C., Mulet, K., and Castagnone-Sereno, P. (2012) The map-1 gene family in root-knot nematodes, Meloidogyne spp.: a set of taxonomically restricted genes specific to clonal species. PloS one, 7, e38656.

Toruño, T.Y., Stergiopoulos, I., and Coaker, G. (2016) Plant-Pathogen Effectors: Cellular Probes Interfering with Plant Defenses in Spatial and Temporal Manners. Annual Review of Phytopathology, 54, 419–441.

Truong, N.M., Nguyen, C.-N., Abad, P., Quentin, M., and Favery, B. (2015) Function of Root-Knot Nematode Effectors and Their Targets in Plant Parasitism. In: Elsevier (Escobar, C. and Fenoll, C., eds), pp. 293–324. Academic Press.

Velasquez, A., Chakravarthy, S., and Martin, G.B. (2009) Virus-induced Gene Silencing (VIGS) in Nicotiana benthamiana and Tomato. Journal of Visualized Experiments, 28: 1292. doi: 10.3791/1292.

Wang, X., Xue, B., Dai, J., Qin, X., Liu, L., Chi, Y., et al. (2018) A novel Meloidogyne incognita chorismate mutase effector suppresses plant immunity by manipulating the salicylic acid pathway and functions mainly during the early stages of nematode parasitism. Plant Pathology, 67, 1436–1448.

Yamaguchi, Y.L., Suzuki, R., Cabrera, J., Nakagami, S., Sagara, T., Ejima, C., et al. (2017) Root-Knot and Cyst Nematodes Activate Procambium-Associated Genes in Arabidopsis Roots. Frontiers in Plant Science, 8, 1195.

Yang, Y., Jittayasothorn, Y., Chronis, D., Wang, X., Cousins, P., and Zhong, G.-Y. (2013) Molecular characteristics and efficacy of 16D10 siRNAs in inhibiting root-knot nematode infection in transgenic grape hairy roots. PloS one, 8, e69463.

Zhang, L., Davies, L.J., and Elling, A. a (2015) A Meloidogyne incognita effector is imported into the nucleus and exhibits transcriptional activation activity in planta. Molecular Plant Pathology, 16, 48–60.

