## Supporting information for "Silencing *SmD1* in Solanaceae alters susceptibility to root-knot nematodes"

>MiEFF18a_312 aa_Minc3s01525g24488
MYFKAFLILLISPFAAFVVNNARTISNMDEVGDVSATMAEKINNVGEKSNQVLPQGEEGMDDLTEEYDDSDDSDEDEDYDDEEDDIDEDNDANNYDYLNSRIDHMDDIDSFETYEELPHMDNYGKMNEAKQMMP
VMHMNAVENEHYKQLPSIATQPIIEANPIMPPMPVVQPQKMEQAPEAMKKEATPQASQPSSPKAATPKAKTSKPKTELKKPSAKKVQAKKGAAKVAKKDTKKPKDVKKSKDAKAKTAKGKVPKPKGKAAKGKPA
KPAKGKPMKGKPAKATKAGPKKQPAKKGKKSASKKVTTKGGKKH
>MiEFF18b_321 aa_Minc3s06052g39242_reannot
MYFKAFLLLLISLLSAFVVNNARTISNMDEVGDVSATMTEKKNNLGEKSNQVLPQGEEGMDDLTEEYDDSDDSEEDEDNDDEEDDMDEDNDANNYDYLNSRIDHMDDIDSFETYEELPHMDNYGKMNEAKQM
MPVMHMNAGENEHYKQLPSIATQPINEANPIMPPMPVVQPEKMEQAPEAMKTEATPQASQPSSPKTAAPKAKTSKPKTELKKPSAKKVQATKGAAKVTKKDAKKPKDVKKSKDPKAKTAKGKVPKAKAAKGKP
AKGKAAKGKPAKPGKGKPLKGKTAKASKAGSKKQPAKKGGKKSGSKKVTAKGGKKH
>MiEFF18c_316 aa_Minc3s00519g13664
MYFKAFLLLLISLLSAFVVNNARTISNMDEVGDVSATMTEKKNNLGEKSNQVLPQGEEEMDDLTEEYDDSDDSEEDEDNDDEEDDMDEDNDANNYDYLNSRIDHMDDIDSFETYEELPHMDNYGKMNEAKQM
MPVMHMNAGENEHYKQLPSIATQPINEANPIMPPMPVVQPEKMEQAPEAMKTEATPQASQPSSPKTAAPKAKTSKPKTELKKPSAKKVQATKGAAKVTKKDAKKPKDVKKSKDPKAKTAKGKVPKAKAAKGKP
AKGKPAKPGKGKPLKGKTAKASKAGSKKQPAKKGGKKSGSKKVTAKGGKKH
>MjEFF18a_312 aa_Mjav1s01285g014445
MYFKAFLILLISLFAALVVNNARTISNMDEVNDVSATMAEKKNNVGEKSNQVLPQGEEGMDDLTEEYDDSDDSDEDEDYDDEEDDIDEDNDANNYDYLNSRIDHMDDIDSFETYEELPHMDNYGKMNEAKQMM
PVMHMNAVENEHYKQLPSIATQPIIEANPIMPPMPVVQPQKMEQAPEAMKKEATPQASQPSSPKAATPKAKTSKPKTELKKPSAKKVQAKKGAAKVAKKDTKKPKDVKKSKDAKAKTAKGKVPKPKGKAAKGKP
AKPAKGKPMKGKPAKATKAGPKKQPAKKGKKSASKKVTTKGGKKH
>MjEFF18b_321 aa_Mjav1s00582g007829
MYFKAFLLLLISLLSAFVVNNARTISNMDEVGDVSATMTGKKNNLGEKSNQVLPQGEEEMDDLTEEYDDSDDSEEDEDYDDEEDDMDEDNDANNYDYHNSRIDHMDDIDSFETYEELPHMDNYGKMNEAKQM
MPVMHMNAGENEHYKQLPSIATQPINEANPIMPPMPVVQPEKMEQAPEAMKTEATPQASQPSSPKTAAPNAKTSKPKTELKKPSAKKVQATKGAAKVTKKDAKKPKDVKKSKDPKAKTAKGKVPKAKAAKGKP
AKGKAAKGKPAKPGKGKPLKGKTAKASKAGPKKQPAKKGGKKSGSKKVTAKGGKKH
>MjEFF18c_312 aa_Mjav1s26636g092645_reannot
MYFKAFLILLISLFAAFVVNNARTISNMDEVGDVSATMAEKINNVGEKSNQVLPQGEEGIDDLTEEYDDSDDSDEDEDYDDEEDDIDEDNDANNYDYLNSRIDHMDDIDSFETYEELPHMDNYGKMNEAKQMMPV
MHMNAVENEHYKQLPSIATQPIIEANPIMPPMPVVQPQKMEQAPEAMKKEATPQAPQPSSPKAATPKAKTSKPKTEFKKPSAKKVQAKKGAAKVAKKDTKKPKDVKKSKDAKAKTAKGKVPKPKGKAAKGKPA
KPAKGKPMKGKPAKATKAGPKKQPAKKGKKSASKKVTTKGGKKH
>MaEFF18a_312 aa_Mare1s12532g079807
MYFKAFLILLIFLFAAFVVNNARTISNMDEVGDVSATMAEKINNVREKSNQVLPQGEEGMDDLTEKYDDSDDSDEDEDYDDEEDDIDEDNDANNYDYLNSRIDHMDDIDSFETYEELPHMDNYGKMNEAKQMIPV
MHMNAVENEHYKQLPSIATQPIIEANPIMPPMPVVQPQKMEQAPEAMKKEATPQASQPSSPKAATPKAKTSKPKTELKKPSAKKVQAKKGAAKVAKKDTKKPKDVKKSKDAKAKTAKGKVPEPKGKAAKGKPAK
PAKGKPMKGKPAKATKAGPKKQPAKKGKKSASKKVTTKGGKKH
>MaEFF18b_312 aa_Mare1s03418g041270
MYFKAFLILLISLFAALVVNNARTISNMDEVNDVSATMAEKKNNVGEKSNQVLPQGEEGMDDLTEEYDDSDDSDEDEDYDDEEDDIDEDNDANNYDYLNSRIDHMDDIDSFETYEELPHMDNYGKMNEAKQMM
PVMHMNAVENEHYKQLPSIATQPIIEANPIMPPMPVVQPQKMEQAPEAMKKEATPQAPQPSSPKAATPKAKTSKPKTEFKKPSAKKVQAKKGAAKVAKKDTKKPKDVKKSKDAKAKTAKGKVPKPKGKAAKGK
PAKPAKGKPMKGKPAKATKAGPKKQPAKKGKKSASKKVTTKGGKKH
>MaEFF18c_321 aa_Mare1s03464g041635
MYFKAFLLLLISLLSAFVVNNARTISNMDEVGDVSATMTGKKNNLGEKSNQVLPQGEEEMDDLTEEYDDSDDSEEDEDYDDEEDDMDEDNDANNYDYLNSRIDHMDDIDSFETYEELPHMDNYGKMNEAKQM
MPVMHMNAGENEHYKQLPSIATQPINEANPIMPPMPVVQPEKMEQAPEAMKTEATPQASQPSSPKTAAPKAKTSKPKTELKKPSAKKVQATKGAAKVTKKDAKKPKDVKKSKDPKAKTAKGKVPKAKAAKGKP
AKGKAAKGKPAKPGKGKPLKGKTAKASKAGPKKQPAKKGGKKSGSKKVTAKGGKKH
>MaEFF18d_321 aa_Mare1s06733g060172
MYFKAFLLLLISLLSAFVVNNARTISNMDEVGDVSATMTGKKNNLGEKSNQVLPQGEEEMDDLTEEYDDSDDSEEDEDYDDEEDDMDEDNDANNYDYHNSRIDHMDDIDSFETYEELPHMDNYGKMNEAKQM
MPVMHMNAGENEHYKQLPSIATQPINEANPIMPPMPVVQPEKMEQAPEAMKTEATPQASQPSSPKTAAPNAKTSKPKTELKKPSAKKVQATKGAAKVTKKDAKKPKDVKKSKDPKAKTAKGKVPKAKAAKGKP
AKGKAAKGKPAKPGKGKPLKGKTAKASKAGPKKQPAKKGGKKSGSKKVTAKGGKKH
>MfEFF18_312 aa_Mfscf7180000422394.g8872
MYFKAFLILLISPFAAFVVNNARTISNMDEVGDVSATMAEKINNVGEKSNQVLPQGEEGMDDLTEEYDDSDDSDEDEDYDDEEDDIDEDNDANNYDYLNSRIDHMDDIDSFETYEELPHMDNYGKMNEAKQMMP
VMHMNAVENEHYKQLPSIATQPIIEANPIMPPMPVVQPQKMEQAPEAMKTEATPQASQPSSPKAATPKAKTSKPKTELKKPSAKKVQAKKGAAKVAKKDTKKPKDVKKSKDAKAKTAKGKVPKPKGKAAKGKPA
KPAKGKPMKGKPAKATKAGPKKQPAKKGKKSASKKVTTKGGKKH
>MhEff18_310 aa_MhA1_Contig1554.frz3.gene5
MNFKAFLLLSISLFSAFVLNNARSISTNMDEVDDVSALMAEKKNNVEENPNLEVIPQGEEEHDDLTEEDDDLDDSEEDDYEDDDDMDDDDDVNNNYDYLNSRIDNMDDIDSFETYEEVPQMDNANYEKTAEVKQI
MPVMHMSAGENEHYKQIPSVITQPIDEANPIVPPMPVAQQQVLPKEATPPASPPASKKAATTKAKPSKAKKPADKKAQVKPKANTKVAKKGEKKSKGAKTSKVTKKANNAKVKTPKGKVPKVKAGKAAKGKPTK
GKPAKGKAGKSKAAKAGPKKQPAKKGGKKPAGKNGAKAKKH
>MlucEFF18a_312 aa_M. luci contig000032F
MYFKAFLILLISLFAAFVVNNARTISNMDEVGDVSATMAEKINNVGEKSNQVLPQGEEGMDDLTEDYDDSDDSDEDEDYDDEEDDIDEDNDANNYDYLNSRIDHMDDIDSFETYEELPHMDNYGKMNEAKQMMP
VMHINAVENEHYKQLPSIATQPIIEANPIMPPMPVVQPQKMEQAPEAMKKEATPQASQPSSPKAATPKAKTSKPKTELKKPSAKKVQAKKGAAKVAKKDTKKPKDVKKSKDAKAKTAKGKVPKPKGKAAKGKPA
KPAKGKPMKGKPAKATKAGPKKQPAKKGKKSASKKVTTKGGKKH
>MlucEFF18b_321 aa_M. luci contig000025F
MYFKAFLLLLISLLSAFVVNNARTISNMDEVGDVSATMTEKKNNLGEKSNQVLPQGEEEMDDLTEEYDDSDDSEEDEDNDDEEDDMDEDNDANNYDYLNSRIDHMDDIDSFETYEELPHMDNYGKMNEAKQM
MPVMHMNAGENEHYKQLPSIATQPINEANPIMPPMPVVQPEKMEQAPEAMKTEATPQASQPSSPKTAAPKAKTSKPKTELKKPSAKKVQATKGAAKVTKKDAKKPKDVKKSKDPKAKTAKGKVPKAKAAKGKP
AKGKAAKGKPAKPGKGKPLKGKTAKASKAGSKKQPAKKGGKKSGSKKVTAKGGKKH
>MlucEFF18c_321 aa_M. luci contig000026F
MYFKAFLLLLISLLSAFVVNNARTISNMDEVGDVSATMTGKKNNLGEKSNQVLPQGEEEMDDLTEEYDDSDDSEEDEDNDDEEDDMDEDNDANNYDYLNSRIDHMDDIDSFETYEELPHMDNYGKMNEAKQM
MPVMHMNAGENEHYKQLPSIATQPINEANPIMPPMPVVQPEKMEQAPEAMKTEATPQASQPSSPKTAAPKAKTSKPKTELKKPSAKKVQATKGAAKVTKKDAKKPKDVKKSKDPKAKTAKGKVPKAKAAKGKP
AKGKAAKGKPAKPGKGKPLKGKTAKASKAGSKKQPAKKGGKKSGSKKVTAKGGKKH
>MeEFF18a_314 aa
MYFKAFLILLISLFAAFVVNNARTISNMDEVDDVSATMAEKKNNVVEKSNQVLPQVEEGMDDLTEEYDDSDDSEEEDDYDDEEDDMDDDNDVNKNDYPNSRIDHMDDIDSFETYEELPYMENYGKMAEAKQMM
PVMHMNAVENEHYKQLPSIATQPINEANPIVPTMPVVQPEKMEQAPEAMKKEATPQASQPSSPKAAAPKAKTSKPKTEVKKPSAKKVQASKVPKKQPAKKDTKKPKDVKKPKDPKAKTAKGKAPKAKGKAAKS
KAAKPAKGKPTKGKPAKATKAGPKKQPAKKGGKKPANKKATTKGGKKH
>MeEFF18b_316 aa_Ment3s00179g0226491
MYFKAFLLLLISLLAAFVVNNARTISNMDEVDDVSATMAEKKNNVGEKSNQVLPQGEEEMDDLTEEYDDSDDSEEDEDYDDEEDDMDEDNDVNNHDYLNSRIDHMDDIDSFETYEELPHMENYGKMAEAKQMM
PVMHMNAVENEHYKQLPSIATQPINEANPIMPPMPVVQPEKMEQAPEAMKTEATPQASQPSSPKAAAPKAKTSKPKTEVKKPSAKKVQATKGAAKVTKKDAKKPKDVKKSKDPKAKTAKGKVPKAKAAKGKPA
KGKPAKPGKGKPLKGKTAKASKAGSKKQPAKKGGKKPANKKATAKGGKKH

**Figure S1** Identified EFF18 sequences in RKN species. aa, amino acids.

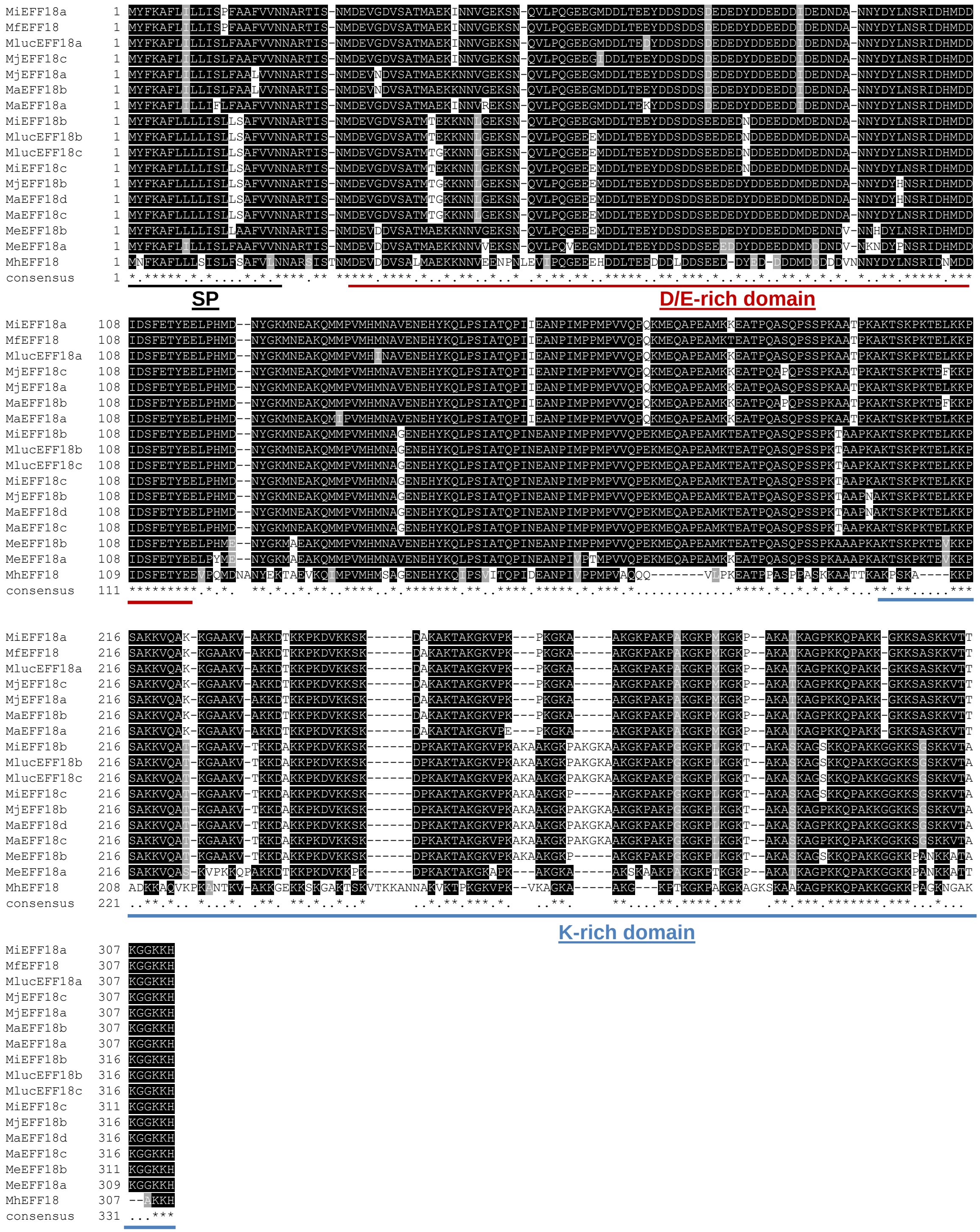


**Figure S2** EFF18 is a conserved RKN effectors. Sequence alignment (MAFFT tool, EBI server) of all identified EFF18 sequences in RKN species. Black line underlines the secretion signal peptide (SP), red line underlines the D/E-rich domain and the blue line underline the K-rich domain. Alignment are colored and visualised using BoxShade.

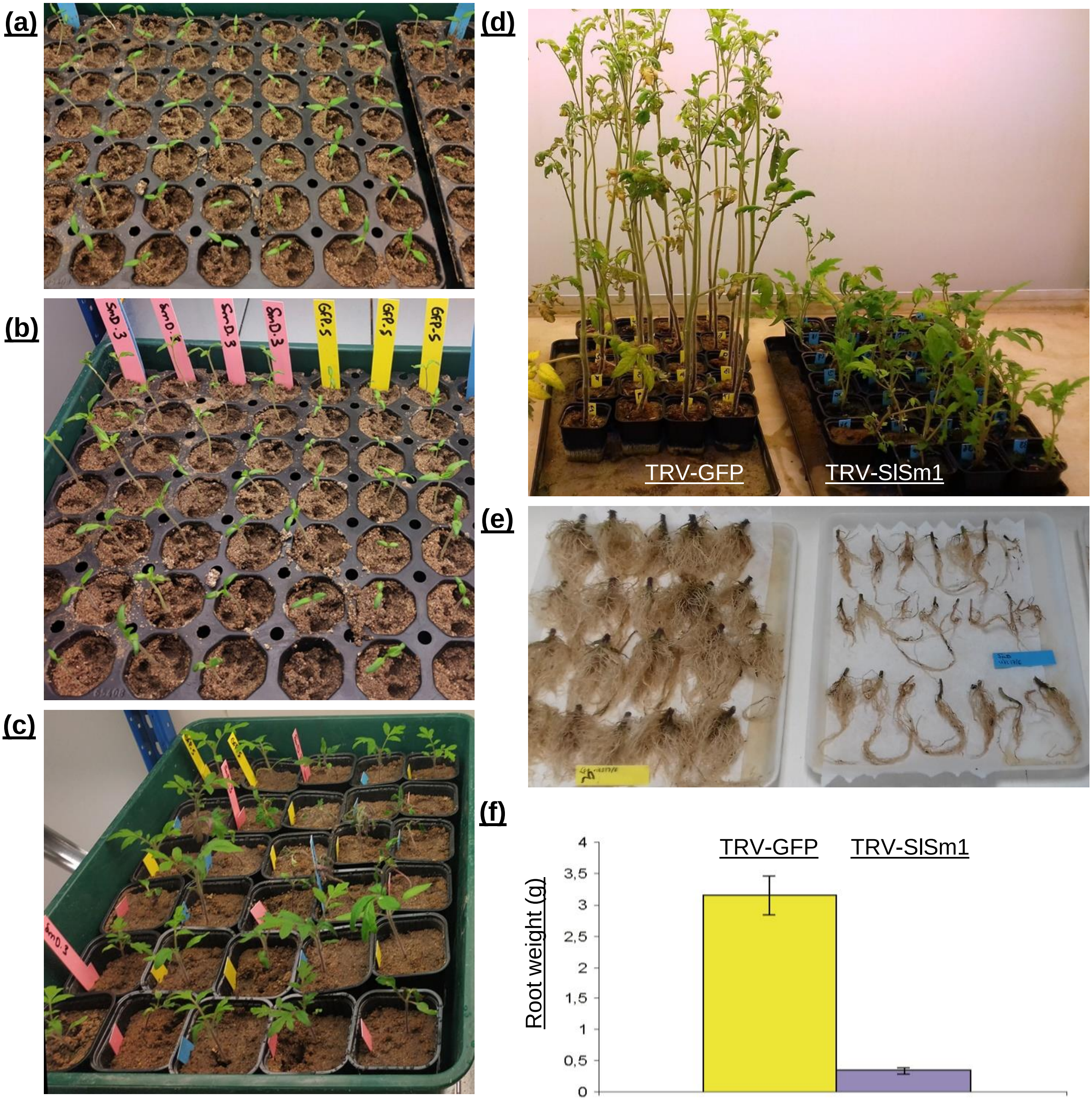


**Figure S3** Tomato phenotypes associated with VIGS of *SlSmD1* genes. (a) 10-day old tomato cotyledons on the day of agroinoculation. (b) Tomato cotyledons 8 days after agroinoculation with TRV1+TRV2-SlSmD1 or TRV2-GFP (VIGS). (c) Tomato cotyledons 4 weeks after VIGS of SlSmD1 (pink labels) or GFP (yellow label) on the day of *M. incognita* inoculation. (d) Shoot phenotype of VIGSed tomato plants, seven weeks after *M. incognita* infection; on the right, TRV-SlSmD1 (blue labels) and on the left, control TRV-GFP (yellow label). (e) Mean root weights of tomato of TRV-SlSmD1 and control TRV-GFP seven weeks after *M. incognita* infection. Error bars represent SEM.

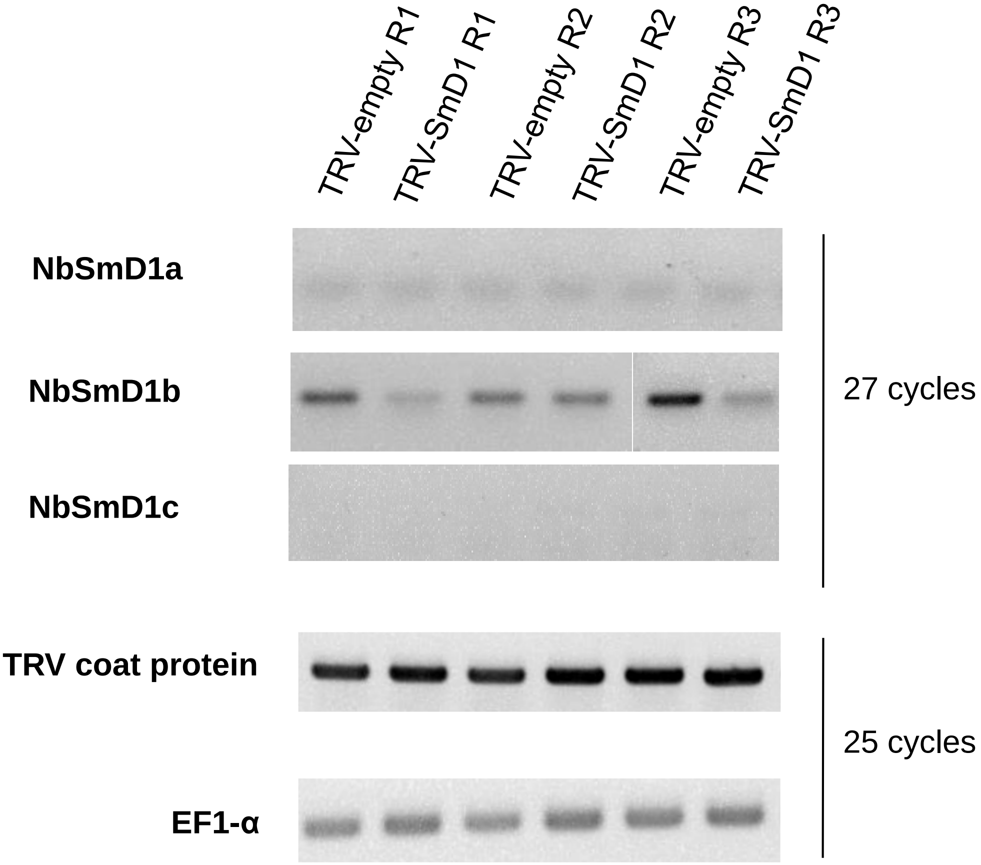


**Figure S4** Semi-quantitative RT-PCR expression analysis of NbSmD1s in *N. benthamiana* VIGS experiments.

### **Table S1** Sequences used in this study and accession numbers

| Species | Gene name | Gene name (from genome) | Accession number | Size (aa) | Signal peptide predicted with Phobius (aa) | NLS mono- and bi-partite (predicted with NUSDISC method, WolfPsort Plant) | Database or references |
|---|---|---|---|---|---|---|---|
| *M. incognita* | MiEFF16 | Minc16401 | MT591035 | 69 | 1 to 21 | pat4: RKHR (3) at 44<br>pat7: PTRKHRP (4) at 42<br>bipartite: none | Ressources INRA (Blanc-Mathieu et al 2017) |
| *M. incognita* | MiEFF18a | Minc18636 / Minc3s01525g24488 | KX907770 | 312 | 1 to 21 | pat4: KKPK (4) at 235<br>pat7: PAKKGKK (4) at 292<br>bipartite: KKGAAKVAKKDTKKPKD at 223 | Ressources INRA (Blanc-Mathieu et al 2017) |
| *M. incognita* | MiEFF18b | Minc3s06052g39242 (reannotated) | | 321 | 1 to 21 | pat4: KKPK (4) at 235<br>pat7: none<br>bipartite: none | Ressources INRA (Blanc-Mathieu et al 2017) |
| *M. incognita* | MiEFF18c | Minc3s00519g13664 | | 316 | 1 to 21 | pat4: KKPK (4) at 235<br>pat7: none<br>bipartite: none | Ressources INRA (Blanc-Mathieu et al 2017) |
| *M. arenaria* | MaEFF18a | Mare1s12532g079807 | | 312 | 1 to 21 | pat4: KKPK (4) at 235<br>pat7: PAKKGKK (4) at 292<br>bipartite: KKGAAKVAKKDTKKPKD at 223 | Ressources INRA (Blanc-Mathieu et al 2017) |
| *M. arenaria* | MaEFF18b | Mare1s03418g041270 | | 312 | 1 to 21 | pat4: KKPK (4) at 235<br>pat7: PAKKGKK (4) at 292<br>bipartite: KKGAAKVAKKDTKKPKD at 223 | Ressources INRA (Blanc-Mathieu et al 2017) |
| *M. arenaria* | MaEFF18c | Mare1s03464g041635 | | 321 | 1 to 21 | pat4: KKPK (4) at 235<br>pat7: none<br>bipartite: none | Ressources INRA (Blanc-Mathieu et al 2017) |
| *M. arenaria* | MaEFF18d | Mare1s06733g060172 | | 321 | 1 to 21 | pat4: KKPK (4) at 235<br>pat7: none<br>bipartite: none | Ressources INRA (Blanc-Mathieu et al 2017) |
| *M. javanica* | MjEFF18a | Mjav1s01285g014445 | | 312 | 1 to 21 | pat4: KKPK (4) at 235<br>pat7: PAKKGKK (4) at 292<br>bipartite: KKGAAKVAKKDTKKPKD at 223 | Ressources INRA (Blanc-Mathieu et al 2017) |
| *M. javanica* | MjEFF18b | Mjav1s00582g007829 | | 321 | 1 to 21 | pat4: KKPK (4) at 235<br>pat7: none<br>bipartite: none | Ressources INRA (Blanc-Mathieu et al 2017) |
| *M. javanica* | MjEFF18c | Mjav1s26636g092645 (reannotated) | | 312 | 1 to 21 | pat4: KKPK (4) at 235<br>pat7: PAKKGKK (4) at 292<br>bipartite: KKGAAKVAKKDTKKPKD at 223 | Ressources INRA (Blanc-Mathieu et al 2017) |
| *M. floridensis* | MfEFF18 | scf7180000422394.g8872 | | 312 | 1 to 21 | pat4: KKPK (4) at 235<br>pat7: PAKKGKK (4) at 292<br>bipartite: KKGAAKVAKKDTKKPKD at 223 | Wormbase parasite (Lunt et al., 2014) |
| *M. enterolobii* | MeEFF18a | | MW272456 | 314 | 1 to 21 | pat4: KKPK (4) at 236; KKPK (4) at 242<br>pat7: none<br>bipartite: KKQPAKKDTKKPKDVKK at 227 | This study |
| *M. enterolobii* | MeEFF18b | Ment3s00179g0226491 | | 316 | 1 to 21 | pat4: KKPK (4) at 236 & KKPK (4) at 242<br>pat7: none<br>bipartite: KKQPAKKDTKKPKDVKK at 227 | (Koutsovoulos et al., 2020) |
| *M. luci* | MlucEFF18a | contig000032F | | 312 | 1 to 21 | pat4: KKPK (4) at 235<br>pat7: PAKKGKK (4) at 292<br>bipartite: KKGAAKVAKKDTKKPKD at 223 | (Susič et al., 2019) |
| *M. luci* | MlucEFF18b | contig000025F | | 321 | 1 to 21 | pat4: KKPK (4) at 235<br>pat7: none<br>bipartite: none | (Susič et al., 2019) |
| *M. luci* | MlucEFF18c | contig000026F | | 321 | 1 to 21 | pat4: KKPK (4) at 235<br>pat7: none<br>bipartite: none | (Susič et al., 2019) |
| *M. hapla* | MhEFF18 | Mhap1s0397g11005 | | 310 | 1 to 24 | pat4: none<br>pat7: PSKAKKP (4) at 201<br>bipartite: KKGGKKPAGKNGAKAKK at 293 | Wormbase parasite (Opperman et al.,2008) |
| *A. thaliana* | AtSmD1a | At3g07590 | Q9SSF1 | 114 | no | pat4: KPKK (4) at 88<br>pat7: PRVKPKK (3) at 85<br>bipartite: KKPVAGKAVGRGRGRGR at 90 | Elvira-Matelot et al., 2016 |
| *A. thaliana* | AtSmD1b | At4g02840 | Q9SY09 | 116 | no | pat4: KPKK (4) at 88<br>pat7: PRIKPKK (3) at 85<br>bipartite: none | Elvira-Matelot et al., 2016 |
| *S. lycopersicum* | SlSmD1a | Solyc06g084310.2.1 | MT598822 | 114 | no | pat4: KPKK (4) at 88<br>pat7: PRVKPKK (3) at 85<br>bipartite: KKPTAGKPMGRGRGRGR at 90 | Mejias et al., 2020 |
| *S. lycopersicum* | SlSmD1b | Solyc09g064660.2.1 | MT598823 | 114 | no | pat4: KPKK (4) at 88<br>pat7: PRVKPKK (3) at 85<br>bipartite: KKPTAGKPMGRGRGRGR at 90 | Mejias et al., 2020 |
| *N. benthamiana* | NbSmD1a | Niben101Scf01782g05006.1 | MT683762 | 114 | no | pat4: KPKK (4) at 88<br>pat7: PRVKPKK (3) at 85<br>bipartite: KKPTAGKPLGRGRGRGR at 90 | This study |
| *N. benthamiana* | NbSmD1b | Niben101Scf05290g01011.1 | MT683763 | 114 | no | pat4: KPKK (4) at 88<br>pat7: PRVKPKK (3) at 85<br>bipartite: KKPTAGKPLGRGRGRGR at 90 | This study |
| *N. benthamiana* | NbSmD1c | Niben101Scf04283g03011.1 | MT683764 | 114 | no | pat4: KPKK (4) at 88<br>pat7: PRVKPKK (3) at 85<br>bipartite: KKPTAGKPLGRGRGRGR at 90 | This study |

**Table S2** Primers used in this study

| Relative to | Name | Sequence (5'- 3') | Purpose |
|---|---|---|---|
| Figure 2 & 3 | pB27_MiEFF18_SfiI_NoATG | GGGGCCGGACGGGCCGCTCGAACCATTTCTAATATG | Cloning bait vector |
| | pB27_MiEFF18_SfiI_STOP | AGGGGCCCCAGTGGCCTTAATGCTTCTTTCCTCC | |
| | pB27_MeEFF18_SfiI_NoATG | GGGGCCGGACGGGCCGCTCGAACCATTTCTAATATG | Cloning bait vector |
| | pB27_MeEFF18_SfiI_STOP | AGGGGCCCCAGTGGCCTTAATGCTTCTTTCCTCCTTTTG | |
| | pB27_MiEFF16_SfiI_NoATG | GGGGCCGGACGGGCCAAGAATAACGACCACCATAAAC | Cloning bait vector |
| | pB27_MiEFF16_SfiI_STOP | AGGGGCCCCAGTGGCCTCATTCATCATCCCCACATTC | |
| | oNP841 | TCAAAACCACTGTCACCT | Sequencing prey vector pP6 |
| | oNP842 | CATAGATCAGGGTTTTCC | |
| | EFF18_F | ATGCCTCCGATGCCAGTTGTCCAACCA | *In situ* hybridisation |
| | EFF18_GW3 | AGAAAGCTGGGTGTTAATGCTTCTTTCCTCCTTTTG | |
| | MeEFF18_GW5 | AAAAAGCAGGCTTCACCATGGCTCGAACCATTTCTAATATG | Gateway cloning entry vector |
| | AtSmD1_GW5 | AAAAAGCAGGCTTCACCATGAAGCTCGTCAGGTTTTTGATG | Gateway cloning entry vector |
| | AtSmD1b_GW3_STOP | AGAAAGCTGGGTGCTAACGACCTCTGCCGCGACCAC | |
| | SlSmD1a_GW5 | AAAAAGCAGGCTTCACCATGAAGCTCGTTAGATTTTTG | Gateway cloning entry vector |
| | SlSmD1a_GW3_STOP | AGAAAGCTGGGTGTTAGCGGCCTCGACCACGTCCAC | |
| | NbSmD1b_GW5 | AAAAAGCAGGCTTCACCATGAAGCTCGTCAGATTTTTG | Gateway cloning entry vector |
| | NbSmD1b_GW3_STOP | AGAAAGCTGGGTGTTAGCGGCCTCGACCACGTCCAC | |
| | AttB1 | GGGGACAAGTTTGTACAAAAAAGCAGGCT | Amplification gateway destination vector |
| | AttB2 | GGGGACCACTTTGTACAAGAAAGCTGGGT | |
| | AttL1 | TCGCGTTAACGCTAGCATGGATCTC | Amplification pDONR207 entry vector |
| | AttL2 | GTAACATCAGAGATTTTGAGACAC | |
| Figure 4 & 5 | TRV2-SlSmD1-F | CAGAATTCAGATTTTTGATGAAGCTGAAC | VIGS |
| | TRV2-SlSmD1-R | GTCTCGAGGACCACGTCCCATAGGCTTTC | |
| | SlSmD1_F1 | GGCCGCTAAATGCATTCCAG | RT-Q-PCR |
| | SlSmD1_R1 | TGCAACGAGATAACATGCTCCT | |
| | SlRPN7_F | TTGGGGTGTCTGAGGATTTC | RT-Q-PCR |
| | SlRPN7_R | CATTCTTTGCATCAGGACGA | |
| | NbSmD1a_F | GGTTGCTGATTGGTGTTATGTC | RT-Q-PCR |
| | NbSmD1a_R | GTAGCTGCATCCACTGAGAG | |
| | NbSmD1b_F | TCGTGTTCGATCTTGCTTTGAC | RT-Q-PCR |
| | NbSmD1b_R | CATCCACACCTGTAATGGTTCC | |
| | NbSmD1c_F | CTCTTTCTACCGCATTCAGTC | RT-Q-PCR |
| | NbSmD1c_R | TCATCACCTTATTTGCTACCCA | |
| | NbACT1_F | GCAACTGGGATGATATGGAG | housekeeping gene RT-Q-PCR |
| | NbACT1_R | TCACGTGTAAGCGAGTTTTTC | |
| | NbEF1α_F | CTGATTATTGACTCCACCACTG | semi quantitative RT-PCR |
| | NbEF1α_R | CATCTTGTTACAGCAGCAAATC | |
| | TRV_CP_F | ACTCACGGGCTAACAGTGCT | semi quantitative RT-PCR |
| | TRV_CP_R | GACGTATCGGACCTCCACTC | |

**References**


Blanc-Mathieu, R., Perfus-Barbeoch, L., Aury, J.-M.M., Da Rocha, M., Gouzy, J., Sallet, E., et al. (2017) Hybridization and polyploidy enable genomic plasticity without sex in the most devastating plant-parasitic nematodes. *PLoS Genetics*, **13**, e1006777.

Elvira-Matelot, E., Bardou, F., Ariel, F., Jauvion, V., Bouteiller, N., Le Masson, I., et al. (2016) The Nuclear Ribonucleoprotein SmD1 Interplays with Splicing, RNA Quality Control, and Posttranscriptional Gene Silencing in Arabidopsis. *The Plant Cell*, **28**, 426–438.

Koutsovoulos, G.D., Poullet, M., Elashry, A., Kozlowski, D.K.L., Sallet, E., Da Rocha, M., et al. (2020) Genome assembly and annotation of Meloidogyne enterolobii, an emerging parthenogenetic root-knot nematode. *Scientific Data*, **7**, 324.

Lunt, D.H., Kumar, S., Koutsovoulos, G., and Blaxter, M.L. (2014) The complex hybrid origins of the root knot nematodes revealed through comparative genomics. *PeerJ*, **2**, e356.

Mejias, J., Bazin, J., Truong, N., Chen, Y., Marteu, N., Bouteiller, N., et al. (2020) The root-knot nematode effector MiEFF18 interacts with the plant core spliceosomal protein SmD1 required for giant cell formation. *New Phytologist*, **in press**, nph.17089.

Opperman, C.H., Bird, D.M., Williamson, V.M., Rokhsar, D.S., Burke, M., Cohn, J., et al. (2008) Sequence and genetic map of Meloidogyne hapla: A compact nematode genome for plant parasitism. *Proc Natl Acad Sci U S A*, **105**, 14802–14807.

Susič, N., Koutsovoulos, G.D., Riccio, C., Danchin, E.G.J., Blaxter, M.L., Lunt, D.H., et al. (2020) Genome sequence of the root-knot nematode Meloidogyne luci. *Journal of Nematology*, **52**, 1–5.
